# The significant yet short-term influence of research covidization on journal citation metrics

**DOI:** 10.1101/2023.03.05.531213

**Authors:** Xiang Zheng, Chaoqun Ni

## Abstract

COVID-19 has emerged as a major research hotspot in recent years, leading to increased publications and citations of related papers. While concerns exist about the potential citation boost in journals publishing these papers, the specifics are not fully understood. This study uses a generalized difference-in-differences approach to examine the impact of publishing COVID-19 papers on journal citation metrics in the Health Sciences fields. Findings indicate that journals publishing COVID-19 papers in 2020 received significantly higher citation premiums due to COVID-19 in 2020 and continued to benefit from the premium in 2021 in certain fields. In contrast, journals that began publishing COVID-19 papers in 2021 experienced weaker citation premiums. Although the publication volume of non-COVID-19 papers also surged, these papers experienced insignificant or negative citation gains, even when published in the same journals as COVID-19 papers. COVID-19 papers published in high-impact journals brought more significant citation premiums to journals in most fields. These citation premiums can affect various citation-based journal metrics, such as our simulated impact factor, to different degrees. The results highlight a “gold rush” pattern in which early entrants establish their citation advantage in research hotspots and caution against using citation-based metrics for research assessment.

## Introduction

Since its appearance in late 2019, the COVID-19 pandemic has caused devastating economic and social disruption. The scientific community is among those who responded swiftly to the widespread and disastrous event (Fraumann & Colavizza, 2022). Scientists have been extensively mobilized and collaborated more than ever to conduct COVID-19-related research to advance knowledge about the virus and develop tools and strategies for controlling the pandemic (Else, 2020; Haghani & Bliemer, 2020; M. Liu et al., 2022; Pai, 2020). As a result, there has been an unprecedented surge in the production of COVID-19 papers, with many active researchers contributing and journals speeding up the publication process (Aviv-Reuven & Rosenfeld, 2021; Else, 2020; Horbach, 2020; Riccaboni & Verginer, 2022; Sarkar et al., 2022): As indexed by Scopus, over 700,000 authors contributed 200,000 COVID-19 papers between January 2020 and August 2021 (Ioannidis et al., 2021). Across disciplines, 98 of the 100 most-cited papers published in 2020-2021 were about COVID-19 (Ioannidis et al., 2022). The surge of COVID-19 papers has led to a massive “covidization” of scientific research during the pandemic, making COVID-19 a significant research hotspot in recent years (Adam, 2020; Ioannidis et al., 2022; Pai, 2020).

The COVID-19 research has indisputably contributed immensely to mitigating and controlling the pandemic, from various protection measures to vaccines and Paxlovid. Nevertheless, researchers in the field expressed concerns that too many scientists pivoting from their professional areas to COVID-19 research could result in tremendous wastage and risk science advances (Pai, 2020). It is also concerned that the massive “covidization” of research will elevate citation-based metrics for certain groups in the research evaluation system, where the citation is widely used as a research impact measure despite widespread criticisms (Aksnes et al., 2019). Researchers and journals that published COVID-19 research would likely have the advantage in the current science evaluation system for publishing COVID-19 research.

Evidence suggests that COVID-19 research brought more citations than non-COVID-19 research to researchers and journals that published them (Brandt et al., 2022; Candal-Pedreira et al., 2022; Ioannidis et al., 2022; W. Liu et al., 2023). It is reported that 98% of the top 100 cited papers published in 2020-2021 were about COVID-19 in Scopus (Ioannidis et al., 2022). Many researchers have achieved citation-based elite status through the disproportionate number of citations received by publishing COVID-19 papers (Ioannidis et al., 2022). For journals, these highly cited COVID-19 papers may seriously boost the journal-level citation count and derivative metrics, such as the Journal Impact Factor (JIF) (Brandt et al., 2022; W. Liu et al., 2023). Studies estimated that journals associated with a high proportion of COVID-19 papers might see increases in JIF from 2020 to 2021 (Fassin, 2021; Gregorio-Chaviano et al., 2022; He et al., 2023; Park et al., 2022; Sjögårde, 2022). Pursuing research hotspots such as COVID-19 appears to yield citation premiums for researchers and journals. However, previous studies have not performed rigorous or large-scale tests to quantify the extent of these citation premiums or the impact on citation-based metrics. Furthermore, most studies only focus on the citation and journal impact trends in the early stages of the COVID-19 pandemic, leaving a research gap in understanding the pandemic’s lasting impact over a longer period.

The COVID-19 pandemic also provides a unique opportunity to unpack the dynamic process of the citation premium brought by research hotspots to journals and researchers. Chasing research hotspots is common in science and can help advance a timely topic and a research field in a short period of time (Wei et al., 2013). It is often assumed that chasing research hotspots can help researchers and journals increase citations and research impact, which is a crucial motivation for such behavior (Bort & Kieser, 2011; Lam, 2011). However, whether research hotspots can consistently produce stable citation premiums is uncertain. Early research suggests that as more papers are published in a field, more citations tend to flow toward established canonical papers rather than new ones (Chu & Evans, 2021). Of the sudden burst of COVID-19 papers, it is possible that late works might not attract as much attention and citations as early works. It remains to be seen whether the COVID-19 trend will continuously benefit journals through citation premiums and how much benefit is left for the “late coming” journals that publish COVID-19 papers later.

Another goal of this study is to examine the mechanism behind the journal citation premium resulting from COVID-19 papers. Highly cited COVID-19 papers directly increased their home journals’ citations (Brandt et al., 2022; Candal-Pedreira et al., 2022; Ioannidis et al., 2022). However, it is unclear whether the timing of publishing COVID-19 papers matters in boosting citations. We are also unsure how COVID-19 papers would influence non-COVID-19 research published in the same journal, given the potential wide visibility brought by COVID-19 papers. While non-COVID-19 papers were published less than before in certain health journals (He et al., 2023) and a citation polarization between COVID-19 and non-COVID-19 papers has been suggested (You et al., 2023), a spillover effect might also occur (Frandsen & Nicolaisen, 2013; Kässi & Westling, 2013): The increased visibility of a journal due to COVID-19 papers may lead to more citations for non-COVID-19 papers in the same journals than would have occurred without the pandemic. Understanding this mechanism will help us comprehend how a research hotspot, directly and indirectly, influences citations and research performance evaluations.

To investigate the research gaps outlined above, this study used a dataset consisting of 5.7 million publications in 7,837 journals indexed by Scopus between 2015 and 2022, including 121,741 COVID-19 publications in 6,303 journals published between 2020-2021, in four *Health Sciences* fields (*Biomedical Research*, *Clinical Medicine*, *Psychology & Cognitive Sciences*, and *Public Health & Health Services*). We quantified the citation premiums brought to journals by COVID-19 papers they published in 2020 and 2021, as well as the resulting changes in common journal impact assessment metrics.

## Data

We obtained research papers published between 2015 and 2022 by journals in the four *Health Sciences* fields from a Scopus data snapshot extracted on March 29, 2024. Journal articles or reviews published in English were included in our analysis. In line with previous research (Brandt et al., 2022; Ioannidis et al., 2021, 2022; Park et al., 2022), we searched the following terms (including different variations with/without hyphens and spaces) in the titles, abstracts, or keywords of all journal papers published from 2020 and onwards to identify COVID-19 papers: *sars-cov-2, coronavirus 2, novel corona, 2019-ncov, ncov-2019, sars-ncov, covid, coronavirus disease 2019, corona-19, corona-2019*, and *hcov-19*. This search retrieved 306,438 unique records across all fields.

We identified the field for each journal based on the journal classification system developed by Science-Metrix (Archambault et al., 2011), which classifies each journal in Scopus into five domains, 20 fields, and 174 subfields. Our analysis focused on four fields within the domain of *Health Sciences*, i.e., *Biomedical Research*, *Clinical Medicine*, *Psychology & Cognitive Sciences*, and *Public Health & Health Services.* These fields were selected due to their direct relevance to COVID-19 and their higher representation of COVID-19 papers and journals publishing them (see **Figure S1**). Furthermore, to pinpoint journals with a multi/interdisciplinary approach that might encompass *Health Sciences* content, we undertook a calculation. Specifically, we determined the proportion of papers labeled as *Health Sciences* papers according to Science-Metrix’s paper-level classification (Rivest et al., 2021), considering all papers published by each journal in 2015-2022. We disregarded the classification at the journal level for this analysis and calculated the share of *Health Sciences* papers in that timeframe for all journals outside *Health Sciences*. Through this methodology, we successfully identified 70 journals that presented over 60 papers in 2015-2022, with 50% or more of their content falling within the category of *Health Sciences* papers. We allocated each of these journals to a subfield that contributed the most to their total paper count and added these journals to our analysis.

We measured the citation impact of a journal using ACPP. To understand the potential impact of COVID-19 papers, we also calculated the COVID-ACPP and the non-COVID-ACPP for these journals (see **Table 1**). We set the last year of ACPP calculation as 2021 and calculated the citation using a three-year time window, which began from the publication year and included that year (van Raan, 2006). We excluded journal self-citations (citations within the same journal) to avoid the potential influence of citation manipulation in the main results. We also replicated the results using total citations (without excluding self-citations) in the robustness check.

**Table 1.** Operationalizations and indications of citation measures.

| Name | Equation | Notation | Indication |
| --- | --- | --- | --- |
| ACPP | $ACPP_{ij} = \frac{\sum_{k=1}^{P_{ij}} C_{ijk}}{P_{ij}}$ | $P_{ij}$: the total paper number of the $i$th journal in year $j$<br>$C_{ijk}$: the number of citations received by the $k$th paper in the $i$th journal in year $j$ | The average citation impact per paper in a journal |
| COVID-ACPP | $COVID - ACPP_{ij} = \frac{\sum_{k=1}^{P'_{ij}} C'_{ijk}}{P'_{ij}}$ | $P'_{ij}$: the total COVID-19 paper number of the $i$th journal in year $j$<br>$C'_{ijk}$: the number of citations received by the $k$th COVID-19 paper in the $i$th journal in year $j$ | The average citation impact of COVID-19 papers in a journal |
| Non-COVID-ACPP | $Non - COVID - ACPP_{ij} = \frac{\sum_{k=1}^{P''_{ij}} C''_{ijk}}{P''_{ij}}$ | $P''_{ij}$: the total non-COVID-19 paper number of the $i$th journal in year $j$<br>$C''_{ijk}$: the number of citations received by the $k$th non-COVID-19 paper in the $i$th journal in year $j$ | The average citation impact of non-COVID-19 papers in a journal |

In addition, we used three citation-based journal metrics to investigate their relationships with the publishing of COVID-19 papers: an impact score simulating the calculation of JIF based on Scopus, SCImago Journal Rank (SJR), and Source Normalized Impact per Paper (SNIP) (Roldan-Valadez et al., 2019). Clarivate proprietarily releases JIF in the *Journal Citation Report* based on the Web of Science database. To simulate JIF in our analysis, we calculated an impact score based on Scopus data for every journal indexed by Scopus using the same methodology:

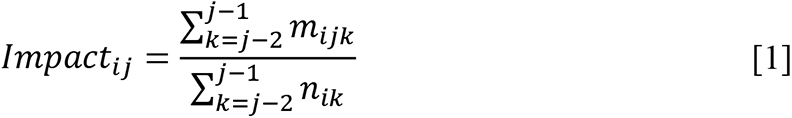

where *Impact_ij_* is the impact score for journal *i* in year *j*, *m_ijk_* is the number of citations made in year *j* to journal *i* ’s past publications published in year *k*, and *n_ik_* is the number of journal *i*’s publications in year *k*. To mitigate the impact of outliers that could skew the results, we further normalized a journal’s impact score to its percentile based on the impact scores of all journals within the same field for a given year and used the impact score percentile in the following calculation. We retrieved the precalculated SJR and SNIP stored in the Scopus dataset directly. Methodologically, SJR considers the prestige of the citing sources and the closeness between cited and citing sources (Guerrero-Bote & Moya-Anegón, 2012), while SNIP normalizes the journal’s citation performance based on the field (Moed, 2010). Both metrics are based on the citations made in the present year to publications in the past three years.

## Methods

### Causal inference strategy

This study used the difference-in-differences (DID) method to evaluate the effect of COVID-19 papers on the citation changes of the entire journal and other non-COVID-19 papers. As a common technique for causal analysis, DID allows us to quantify the influence of publishing COVID-19 research while excluding the impact of other citation inflation factors, such as the annually growing publication volume and the extended reference lists in recent publications (Neff & Olden, 2010; Thelwall & Sud, 2022). We used DID to estimate the impact of COVID-19 research on citations impacts based on the following considerations: If the citation change pattern for journals follows a parallel trend before they published COVID-19 papers, but journals that published COVID-19 papers (the treatment group) deviate from the parallel trend that journals never published COVID-19 papers (the control group) are still following, then the deviation should be caused by publishing COVID-19 papers if no other unobserved events interfere.

### Sample construction

In our dataset, journals that published COVID-19 papers became the treatment group. The rest of the journals became the control group. We performed an exact matching between journals in the treatment and control group to improve the comparability: We subdivided the sample into subclasses by grouping journals based on subfield (the finest level of classification defined by Science-Metrix) and journal open-access (OA) status. Subclasses containing at least one journal in both the treatment and control groups were retained for further analysis. The final sample includes 6,303 journals in the treatment group (publishing 121,741 COVID-19 papers during 2020-2021) and 1,534 “comparable” journals in the control group (see **Table 2**). Due to the potential dynamic treatment effects brought by the time factor (Sun & Abraham, 2021), we divided the treatment group into two cohorts, the 2020 and 2021 cohorts, including journals that started publishing COVID-19 papers in 2020 and 2021, respectively. Comparing each cohort with the control group allows us to examine whether publishing COVID-19 papers brings the same citation premiums over time. Understanding that citation practices vary by field, we constrained all analyses within the same field.

**Table 2.** Number of journals in the sample.

|  |  | *Biomedical*<br>*Research* | *Clinical*<br>*Medicine* | *Psychology &*<br>*Cognitive Sciences* | *Public Health &*<br>*Health Services* |
| --- | --- | --- | --- | --- | --- |
| Treatment<br>Group | 2020-cohort | 569 | 3164 | 175 | 534 |
|  | 2021-cohort | 255 | 1050 | 283 | 273 |
| Control Group |  | 284 | 771 | 317 | 162 |
| Total |  | 1108 | 4985 | 775 | 969 |

### Model specification

We used the generalized DID technique to compare the citation counts of the journals in the treatment and control groups before and after treatment (publishing COVID-19 papers) at different levels of COVID-19 publishing intensity (Kraus & Koch, 2021; Nunn & Qian, 2011). The COVID-19 publishing intensity is measured by the percentage of COVID-19 papers relative to the total number of papers for a journal in a given year, ranging from 0 to 100 (You et al., 2023). It equals 0 for the years before publishing COVID-19 research. We selected the five years before the COVID-19 pandemic (2015-2019) as the pre-treatment period, and 2020-2021 as the in-treatment period. We estimated the following fixed-effect regression specification as the baseline specification:

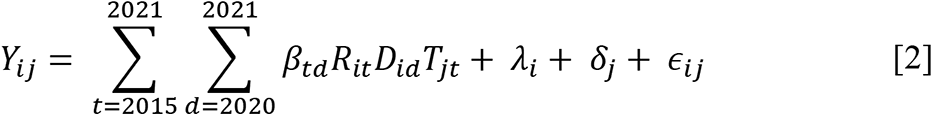

where *i*, and *j* denote the *i*th individual journal and *j*th year, respectively. *Y* denotes the outcome variables of interest, including the ACPP and the non-COVID ACPP. *R_it_* is the treatment variable, the percentage of COVID-19 research in the *i*th individual journal for year *t*. *D_id_* is a cohort dummy variable that equals 1 if the *i*th individual journal starts publishing COVID-19 papers in the year of *d*, belonging to the *d*-cohort. *T_jt_* is a dummy year variable that equals to 1 if *j* = *t*. *λ* and *δ* denote the journal and year-fixed effects, respectively, capturing unobservable journal- and year-specific factors, such as the journal field, specialty, OA status, and yearly changes of citing sources.

We are interested in the coefficients *β_td_*, the estimated differences per 1% of COVID-19 research in journals between the *d*-cohort and the reference group, i.e., journals that never published COVID-19 research in year *t* . *β*_2020,2020_ and *β*_2021,2021_ measure the impact of publishing COVID-19 research on ACPP in 2020 and 2021, respectively, under the assumption of the pre-treatment parallel trend. *β*_2021,2020_ represents the 2020 cohort’s advantage in ACPP in 2021 after publishing COVID-19 research in 2020. Due to the multiple periods for individual journals, we clustered the standard errors at the individual journal level.

Using DID requires the parallel trend assumption that the trends of ACPP over time for each cohort in the treatment group and the control group before COVID-19 are the same. We compared the trends of ACPP for journals with different percentages of COVID-19 research by estimating the following specifications.

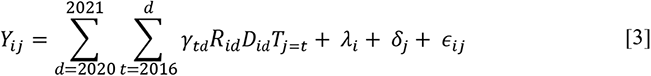

where the coefficient *γ_td_* for the interaction terms of dummy cohort variables and dummy year variables is the estimated differences per 1% of COVID-19 papers between the *d*-cohort and the control group in year *t*. If *γ_td_* for the periods before the first-year publishing COVID-19 papers is not significantly different from 0, we determine that the compared journals with different percentages of COVID-19 publications in corresponding cohort years would have followed a similar trend of ACPP if COVID-19 had not happened. We dropped the starting year of our study (2015) from the interaction terms to avoid multi-collinearity. It should be noted that the generalized DID models with continuous treatment variables may require a stronger assumption of parallel trends than DID models with binary treatment variables (Callaway et al., 2021). We hence converted the treatment variable into a binary variable and examined the parallel trends again in the robustness check (see Robustness check section).

In addition, we checked whether the citation effect from COVID-19 research depends on journal features using heterogeneity analysis. For journal features, we assigned journals into groups based on journal past impact (high- and low-impact journals) and OA status (OA and non-OA journals). *X*% high-impact journals are defined as journals whose 2020 impact score (the latest year not affected by COVID-19) is in the top *X*% of their affiliated fields, and low-impact journals include the rest. We chose the year 2020 because it supposedly captures journal impact rank without potential citation inflation boosted by COVID-19 research. The heterogeneity test estimated the following specifications.

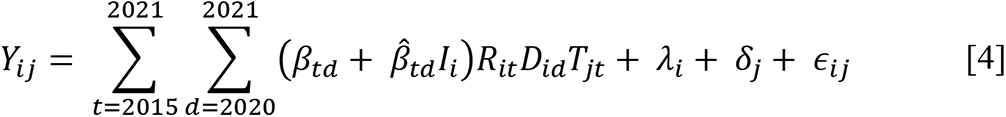

Where *I_i_* is a binary variable denoting the *i*th journal’s group by impact rank and OA status. *β̂_td_* shows the heterogeneous effect between two journal groups according to the definition of *I_i_*.

## Results

### The covidization of scientific research

After 2020, the scientific community has seen a marked increase in research focusing on COVID-19. Between 2020 and 2022, across all *Health Sciences* fields, there was a steady rise in the proportion of COVID-19 papers and journals publishing COVID-19-related research, followed by various degrees of decline in 2023 (see **Figure 1A** and **B**). The average citations of COVID-19 papers, relative to all same-field papers, have declined since 2020 (see **Figure 1C**). This trend suggests that their citation impact may be waning despite the increasing volume of COVID-19 publications. Moreover, *Health Sciences* journals have been publishing a greater number of papers since 2020 than expected, based on the publishing trends of the pre-COVID-19 era (see **Figure 1D**). The publication numbers for non-COVID-19 papers have also surpassed expected levels, indicating that the pandemic has similarly boosted their publication.

**Figure 1.**
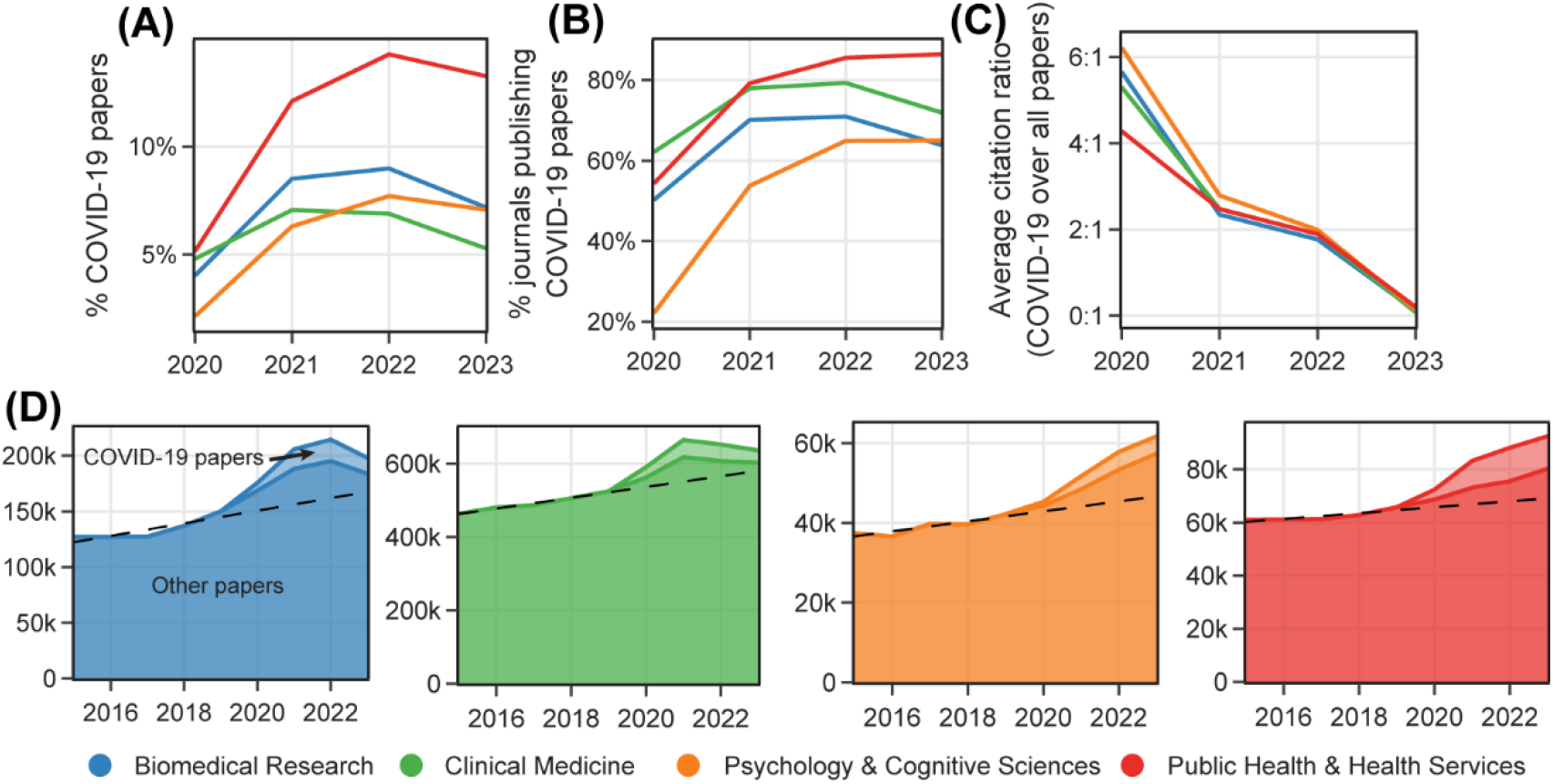
Changes in papers, journals, and citations associated with COVID-19 research. *Note.* (A) Percentages of COVID-19 research papers out of all research papers in the same field. (B) Percentages of journals publishing COVID-19 papers out of all research journals in the same field. (C) Ratios between the average citations of COVID-19 papers and all research papers in the same field. 2022 and 2023 use 2-year and 1-year citation windows, respectively. (D) Changes in total number of papers in *Health Sciences* fields. Dashed lines represent the OLS regression lines fitted from the total number of papers published between 2015 and 2019 (the pre-COVID-19 era), to forecast the expected total number of papers after 2020.

### Trends of citation impact

Figure 2A illustrates the evolution of citation impact for different journal groups since 2015. We see that the average citation per paper in a journal (ACPP) exhibits a slow and stable increase in parallel trends across journal groups in all four fields before 2020. However, potential citation inflation for journals occurred due to publishing COVID-19 papers: The 2020-cohort journals experienced a significant rise by around 28.2% (*Clinical Medicine*)–58.5% (*Psychology & Cognitive Sciences*) in the ACPP from 2019 to 2020. We listed the top five journals with the largest increases in ACPP for each field in **Table S1**. On the other hand, the 2021-cohort journals show a limited deviation in citation measures from the control group for papers published in 2021. We also plotted the evolution of ACPP, similar to Figure 1A, for every other field. We noticed that in some fields, such as *Social Sciences* and *Economics & Business*, the ACPP trends for the defined journal groups are similar to Health Sciences fields, which suggests that COVID-19 might also play a role in these fields’ citation patterns (see **Figure S2**).

**Figure 2.**
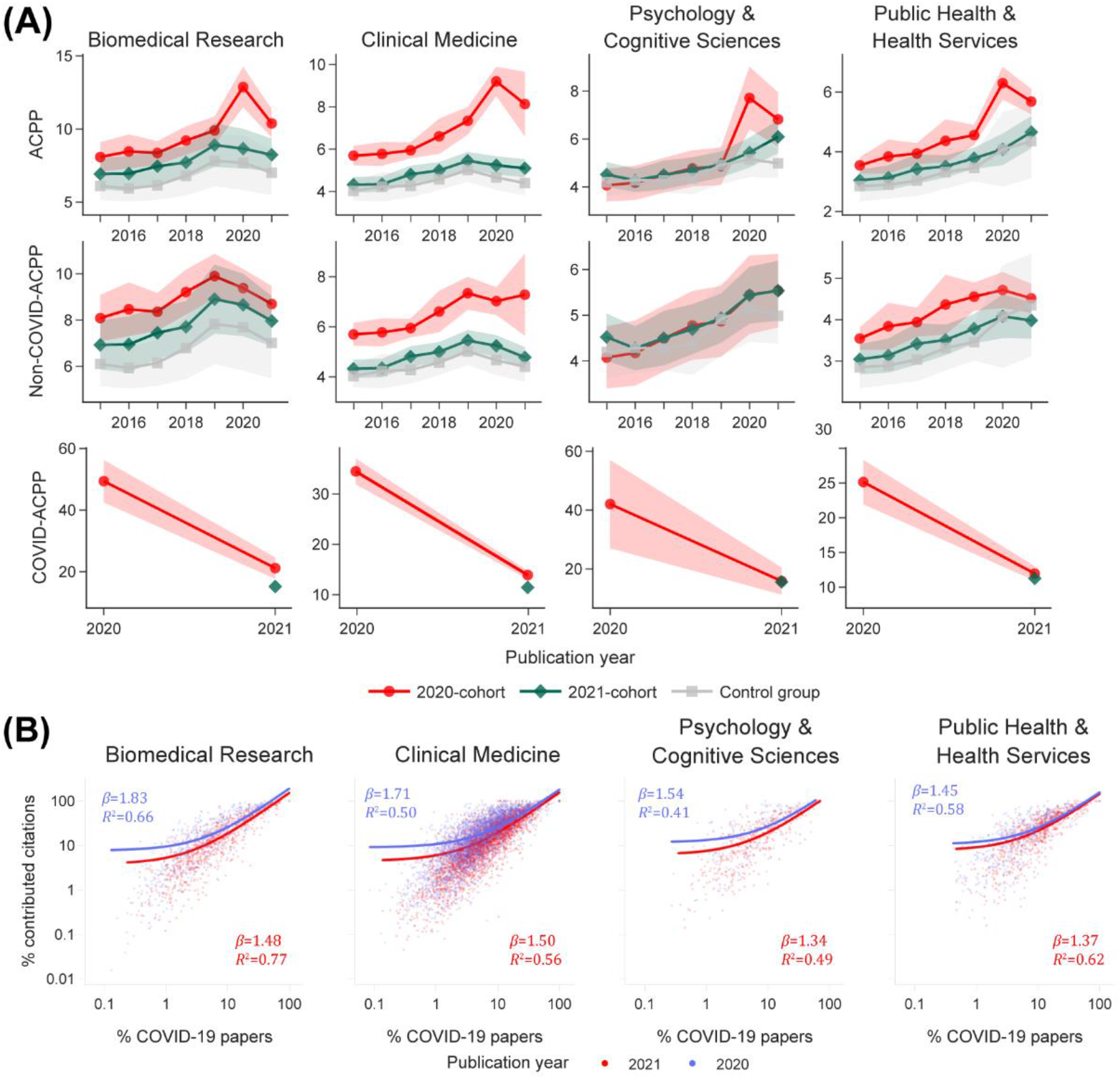
The citation impact of journals by COVID-19 publishing status. *Note.* All citations were calculated using a three-year time window. (A) The evolution of ACPP, non-COVID-ACPP, and COVID-ACPP from 2015-2021. The markers show the mean values of the year. The shaded areas show 95% confidence intervals. (B) The relationship between the percentage of COVID-19 papers in a journal and the percentage of citations contributed by COVID-19 papers during a three-year time window. Trend lines were fitted using OLS. Both horizontal and vertical axes are on the log scale.

The sharp increase in journal citation measures is highly likely due to the COVID-19 papers they published. As the third row in Figure 2A shows, on average, COVID-19 papers published in 2020 have attracted remarkably more citations than non-COVID-19 papers published in the same year. For example, in *Biomedical Research*, the average citation per COVID-19 paper in a journal (COVID-ACPP) (mean=49.30, 95%CI [42.44, 56.16]) in the 2020 cohort is 4.3 times the concurrent ACPP (mean=11.47, 95%CI [11.47, 14.23]) and 5.27 times the non-COVID-ACPP (mean=9.36, 95%CI [8.54, 10.18]) in the same journals. However, the COVID-ACPP declined in 2021 by 52.6% (*Public Health & Health Services*)–62.3% (*Psychology & Cognitive Sciences*), suggesting that COVID-19 papers’ citation impact could quickly shrink. As Figure 2B shows, a small percentage of COVID-19 papers contributed significantly to their home journals’ overall citations. However, COVID-19 papers published in 2020 contributed more to their home journals’ citation measures than those published in 2021, as suggested by the OLS fitting slopes (*β*) of the fitted trend lines. It indicates that the high citation peaks due to COVID-19 papers published in 2020 did not extend to those published in 2021.

We also considered journals’ varying degrees of covidization and grouped the journals by their percentages of COVID-19 research. We segregated the journals from the 2020 and 2021 cohorts into three distinct groups using thresholds of 5% and 10% for the COVID-19 paper percentage (see **Figure S3** for the distribution of COVID-19 paper percentages across different fields). Notably, within the 2020 cohort, journal groups with higher COVID paper percentages exhibit more pronounced surges in ACPP after 2020 (see **Figure S4A**), while the groups within the 2021 cohort generally maintain a trajectory close to the control group (see **Figure S4B**). This observation further lends support to the claim that the citation boosts related to the COVID-19 paper percentages.

### Citation premium due to COVID-19 papers

DID allows us to quantify the substantial citation premium for journals that published COVID-19 papers. As Figure 3A shows, compared with journals in the control group, a 1% increase in COVID-19 papers in 2020-cohort journals on average leads to about 0.36 (95%CI [0.22, 0.49]) in *Biomedical Research,* 0.23 (95%CI [0.18, 0.29]) more ACPP in *Clinical Medicine*, 0.17 (95%CI [0.10, 0.24]) in *Psychology & Cognitive Sciences*, and 0.11 (95%CI [0.07, 0.15]) in *Public Health & Health Services*. Most 2020-cohort journals (91.6%) continued to publish COVID-19 papers in 2021, which brought minor citation advantages to those journals in certain fields. The ACPP of 2021-cohort journals makes less difference than the control group, suggesting a weaker citation premium brought to 2021-cohort journals.

**Figure 3.**
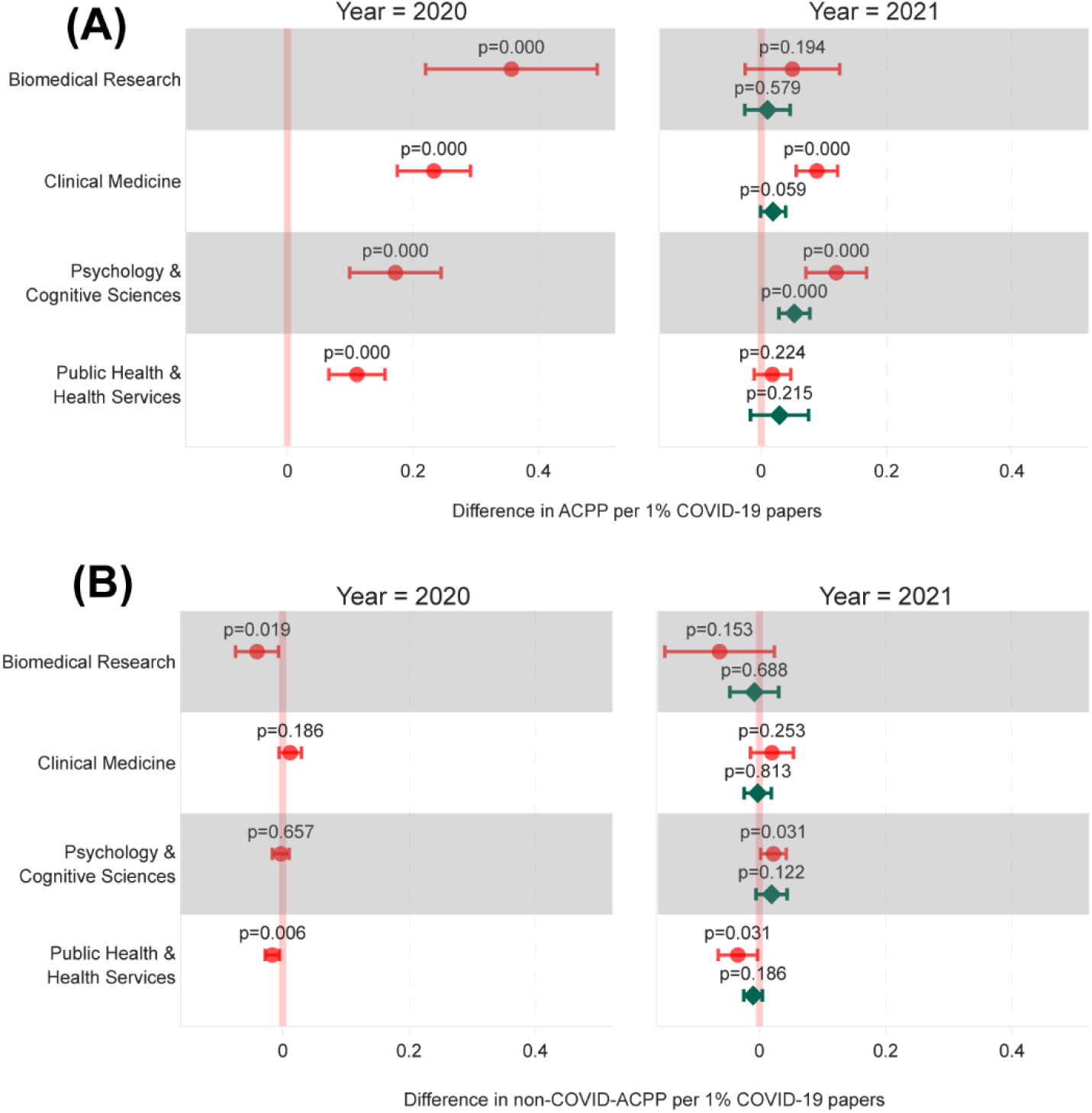
Quantification of journal citation premiums due to COVID-19 papers. *Note.* See **Methods** for regression specifications. Horizontal error bars indicate the 95% confidence intervals. From top to bottom, the numbers of observations by field are 7239, 31268, 4979, and 6143. (A) Effect of publishing COVID-19 papers on ACPP. (B) Effect of publishing COVID-19 papers on non-COVID-ACPP.

The results also suggested that publishing COVID-19 papers did not help increase and might even lower the citations for non-COVID-19 papers in the same journals (See Figure 3B). Compared with journals in the control group, the increase in COVID-19 papers is not shown to attract more citations for their non-COVID-19 papers in *Clinical Medicine* and *Psychology & Cognitive Sciences* and appears to lead to a slight decrease in the other two fields. This holds for both 2020- and 2021-cohorts and both years.

### Robustness check

We confirmed that the main results mentioned above are robust and insensitive to certain factors based on additional robustness tests, including an in-time placebo test, an in-space placebo test, a balanced panel test, an ACPP alternative measure-based test, a binary treatment-based DID test, and additional tests with control variables.

#### Parallel trend test

To formally test the pre-treatment parallel trend assumption required by DID, we used event-study analysis and confirmed the pre-treatment parallel trend. Figure 4 shows that the ACCP difference (coefficient) between the 2020 cohort and the control group was insignificant for research published for most years in 2015-2019. However, the ACPP difference between the 2020 cohort and the control group became significant for research published in 2020 in all four fields. The ACPP difference between the 2021 cohort and the control group remains insignificant for research published between 2015 and 2021. We observed similar patterns regarding non-COVID-ACPP. These results attest to the parallel trend assumption in our analytical sample and the existence of citation boosting effect by COVID-19 papers published in 2020.

**Figure 4.**
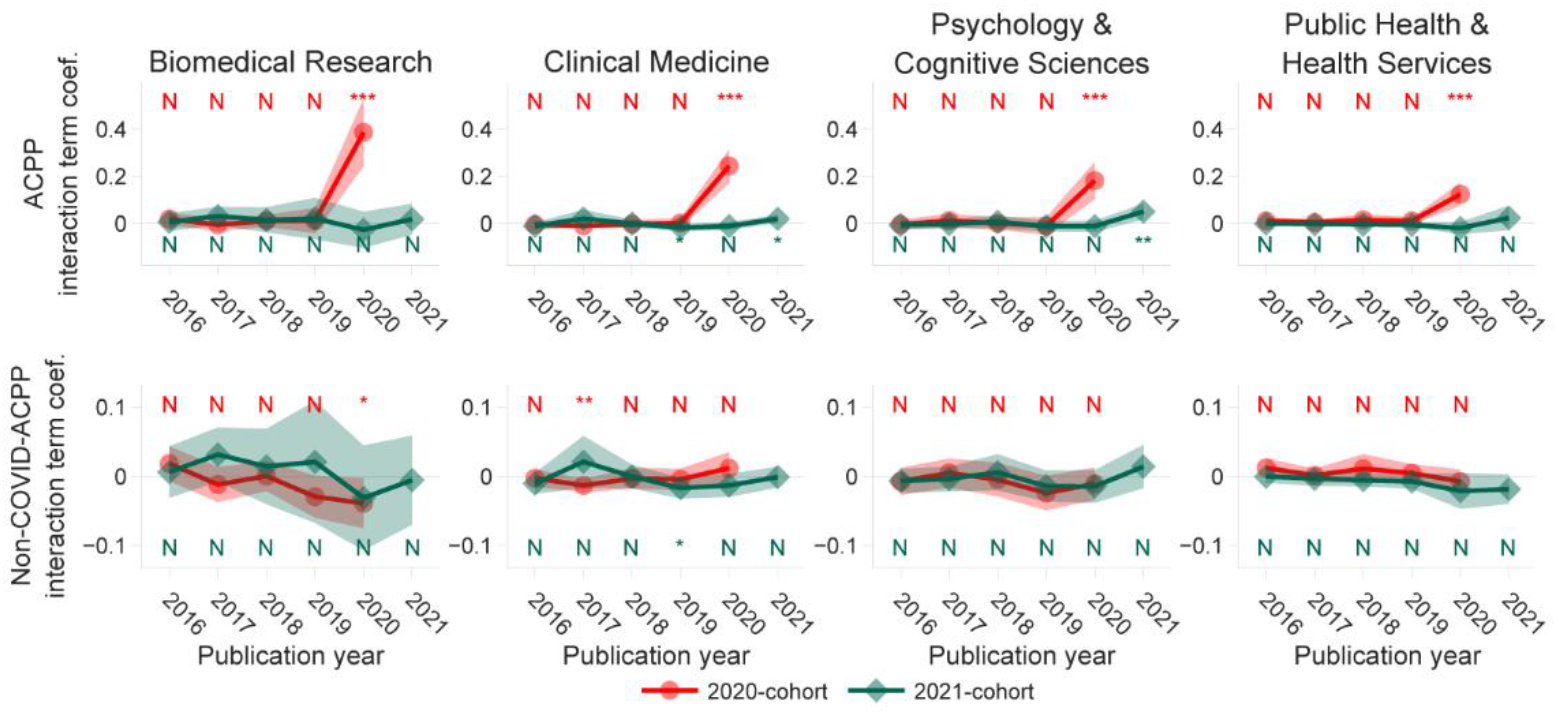
Parallel trend test. *Note.* The y-axis and the markers show the coefficients denoted as ***γ*** in the specification, indicating the estimated differences regarding the outcome variable between the corresponding cohort and the control group in a certain year. The shaded areas show 95% confidence intervals. 2020-cohort’s significance is shown in red at the top; 2021-cohort’s significance is shown in green at the bottom. *** p<0.001, ** p<0.01, * p<0.05, N not significant.

#### Potential confounding events

It is not unlikely that the sharp increase in the 2020 ACPP for journals in the 2020 cohort was because of other unobservable confounding events other than the COVID-19 papers they published. Sudden changes in other citation-influencing factors after 2020 might be a potential cause of the observed citation deviation from the parallel trend. To test this possibility, we delved into the recent temporal patterns regarding the average length of titles, abstracts, number of authors, and references within the timeframe of 2015 to 2021. **Figure S5** illustrates our findings, revealing that there were no such abrupt shifts occurring after the year 2020. Based on this evidence, we maintain the validity of the established causality.

#### Placebo tests

##### In-time placebo test

Beyond the selected confounding factors, we also conducted in-time and in-space placebo tests to examine if other unobservable confounding events interfered with the results. In the in-time placebo test, we assume that the treatment journal group published COVID-19 papers two years before the actual event, i.e., 2018-2019, a period close to the event year but without the intervention of COVID-19 papers. If other unobservable confounding events were present during this period, we would expect significant ACPP rises among journals in the treatment group in the “placebo” treatment years. Our tests show that none of the differences between the treatment group and the control groups is significantly different from zero in the placebo treatment years, suggesting that confounding events are improbable in this study (see **Figure S6**).

##### In-space placebo test

The in-space placebo test intends to rule out unobserved event bias and reverse causality (Hu et al., 2022). We use this test to increase our confidence that the causality between publishing COVID-19 papers and citation increases is true. We randomly permuted the COVID-19 publication records among journals within the same subfield for 2020-2021, generating a shuffled set of treatment and control groups. We then estimated the coefficients of our baseline regression specification using this altered dataset across four fields. We repeated the process 1000 times to generate empirical three-tailed *p*-values, which shows how likely the permuted treatment groups have the same or more significant effect on the outcome variable than the real treatment group. **Table S2** shows that for every significant difference detected between the actual treatment and control group in Figure 3, the empirical p-values in this placebo test remain significant, indicating the actual effects of COVID-19 on ACPP are not achieved by random. All journals in the 2021 cohort’s empirical *p*-values are insignificant (>=0.05), which also aligns with the results based on the actual treatment and control group. In sum, the two placebo tests reduce the possibility of unobserved confounding events impacting our previous findings to a large extent, suggesting robustness in our conclusions.

#### Balanced panel construction

To increase the statistical power, we used all available data in our previous DID tests, which constructed an unbalanced panel dataset. One potential risk introduced by this is that our estimates may not involve all journals during each time point. For instance, journals established in 2018 were not included in the estimates of parallel trends prior to the year. To rule out the potential noise an unbalanced panel may bring to our DID inferences, we converted our dataset into a balanced panel by excluding 1,604 journals that started publishing after 2015 or had missing years. We then examined the parallel trends based on the balanced panel data and re-estimated our specifications to check for the robustness of our previous findings. We found that the new estimates based on the balanced panel data remain unchanged, suggesting minimal biases introduced by the unbalanced panel data used in our findings (see **Figure S7**).

#### Alternative outcome variable based on full citation counts

In the previous DID estimates, we removed journal self-citations (the citing and cited paper are from the same journal) in calculating various citation measures. Ongoing debates exist on the inclusion of journal self-citations in measuring their citation impact (Siler & Larivière, 2022). In light of the debate, we substituted the outcome variables with new ACPP values based on the full citation counts (without excluding self-citations) to test the robustness of our previous findings. Our estimates based on journal full citation counts show limited deviations from the previous results (see **Figure S8**), suggesting our results are insensitive to journal self-citations.

#### Using binary treatment variable

Instead of the generalized DID, we dichotomized the percentage of COVID-19 papers into a binary dummy variable to run standard DID and parallel trend tests with the reclassified treatment and control groups. We use two cutoffs, 0% and 5%, respectively, to dichotomize the percentages of COVID-19 papers. Specifically, all percentages of COVID-19 papers greater than the selected cutoff will be assigned the value of 1, with other rates remaining the same as 0. As shown in **Figure S9,** the estimated results using either of the cutoffs support our conclusions drawn from the main results.

#### Additional Control variables

Although the time-invariant variables are controlled by fixed effects, the uncontrolled time-variant variables may influence the estimates. We examined whether our previous findings are subject to two journal-level time-variant variables, the published volume of journals and the journal impact, which may impact the likelihood of the publications being cited. We used the impact score to proximate journal impact. To avoid being drastically affected by some outliers of highly cited papers, we converted the raw impact score into percentiles among all same-field journals’ impact scores in the same year. Then, we ran the regression using each year’s impact score percentile and publication volume as the control variable. As suggested by **Figure S10**, controlling for these variables does not notably change our estimated results, further confirming our previous findings’ robustness. Using SJR or SNIP percentiles as the proxy of journal impact generates similar results.

### Heterogeneity analysis

After finding COVID-19 research’s significant effect on ACPP, we ask if the effect differs between journals with different features. Journal rank and accessibility are two major journal-level factors that can potentially affect their citations (Tahamtan et al., 2016). We conducted a heterogeneity test to examine if the COVID-19 papers’ citation premium differs across these features. We split journals into subgroup pairs for heterogeneity analysis based on their impact rank and OA status. We first created two pairs of journal subgroups based on journal impact rank. The first pair includes a set of journals within the top 25% impact rank measures (25% high-impact) and 75% impact rank measures (75% low-impact). The second pair includes 50% high- and 50% low-impact journal groups. We also created a journal group pair based on their OA status: OA and non-OA journal subgroups.

To test, we quantified the effect gaps per 1% increase of COVID-19 papers between each pair of journal subgroups (see **Table 3**; complete details in **Table S3**). We found that publishing COVID-19 papers in 2020 increased the journals’ citations more for high-impact journals than for low-impact journals in all fields. Illustrations of the correlation between the journal impact and the ACPP per 1% COVID-19 papers further show that they are positively correlated (see **Figure S11**). For both cohorts, we observed significant positive citation effects for high-impact journals in 2021 in most fields, although the 2021-cohort journals have weaker heterogeneous effects from publishing COVID-19 papers in 2021. Additionally, OA journals’ citation counts were boosted more than non-OA journals’ citation counts by COVID-19 papers published in 2020 in *Clinical Medicine*. We also replicated the same specifications used in the main results to determine if any subgroup differed significantly from the main findings. Our analysis demonstrated that the estimated results were qualitatively similar to the previous main results, strengthening the validity of our estimations for journals with varying levels of impact rank and accessibility (see **Figure S12-S14**).

**Table 3.** Heterogeneity analysis of COVID-19 research effects on ACPP by journal impact rank and OA status.

| Journal cohort | Publication year | Biomedical Research | Clinical Medicine | Psychology & Cognitive Sciences | Public Health & Health Services |
| --- | --- | --- | --- | --- | --- |
| **25% high vs. 75% low-impact** |  |  |  |  |  |
| 2020 | 2020 | 0.801*** | 0.741*** | 0.582*** | 0.302*** |
|  | 2021 | -0.068 | 0.220* | 0.268** | -0.023 |
| 2021 | 2021 | -0.360* | -0.012 | 0.089 | 0.074 |
| **50% high vs. 50% low-impact** |  |  |  |  |  |
| 2020 | 2020 | 0.537*** | 0.507*** | 0.417*** | 0.260*** |
|  | 2021 | -0.022 | 0.138* | 0.212*** | 0.021 |
| 2021 | 2021 | -0.15 | 0.009 | 0.076* | 0.068 |
| **OA vs. subscription-based** |  |  |  |  |  |
| 2020 | 2020 | -0.18 | -0.101* | 0.032 | 0.084* |
|  | 2021 | 0.039 | -0.06 | 0.032 | 0.008 |
| 2021 | 2021 | 0.055 | 0.006 | -0.011 | -0.048 |

**Note.** Numbers in cells are interaction-term coefficients of regression and are proportional to the cell color scale. Only significant coefficients are colored. Green – positive coefficients. Red – negative coefficients. \*\*\* $p < 0.001$ , \*\* $p < 0.01$ , \* $p < 0.05$ .

### Impact on journal metrics

Hypothetically, the citation boost brought by COVID-19 papers should ripple across various citation-based journal metrics. To examine the potential impact, we presented an impact score percentile based on Scopus journals in our dataset (see **Data**). We investigated if the impact score percentiles of journals within the same field would be swayed by COVID-19 in 2021-2022 when the impact score calculation first included the citations to 2020 publications. As Figure 5A shows, 2020-cohort journals in all fields saw an increase in impact score percentiles after 2021. Journals that published a high share of COVID-19 papers tended to have significant increases in rank (see Figure 5B). Additionally, we incorporated alternative journal metrics like SJR and SNIP into our analysis. These alternative metrics utilize more intricate weighting algorithms and longer time frames, rendering them less susceptible to the short-term citation surge resulting from the COVID-19 pandemic’s influence on citations within a single year. Despite these measures, we still discern a subtle effect (see **Figure S15)**. Our findings suggest that, at least in the early stage, citation-based journal metrics are likely to be impacted by the previously suggested COVID-19 premium for 2020-cohort journals, but better-normalized metrics may demonstrate greater robustness against such anomalies.

**Figure 5.**
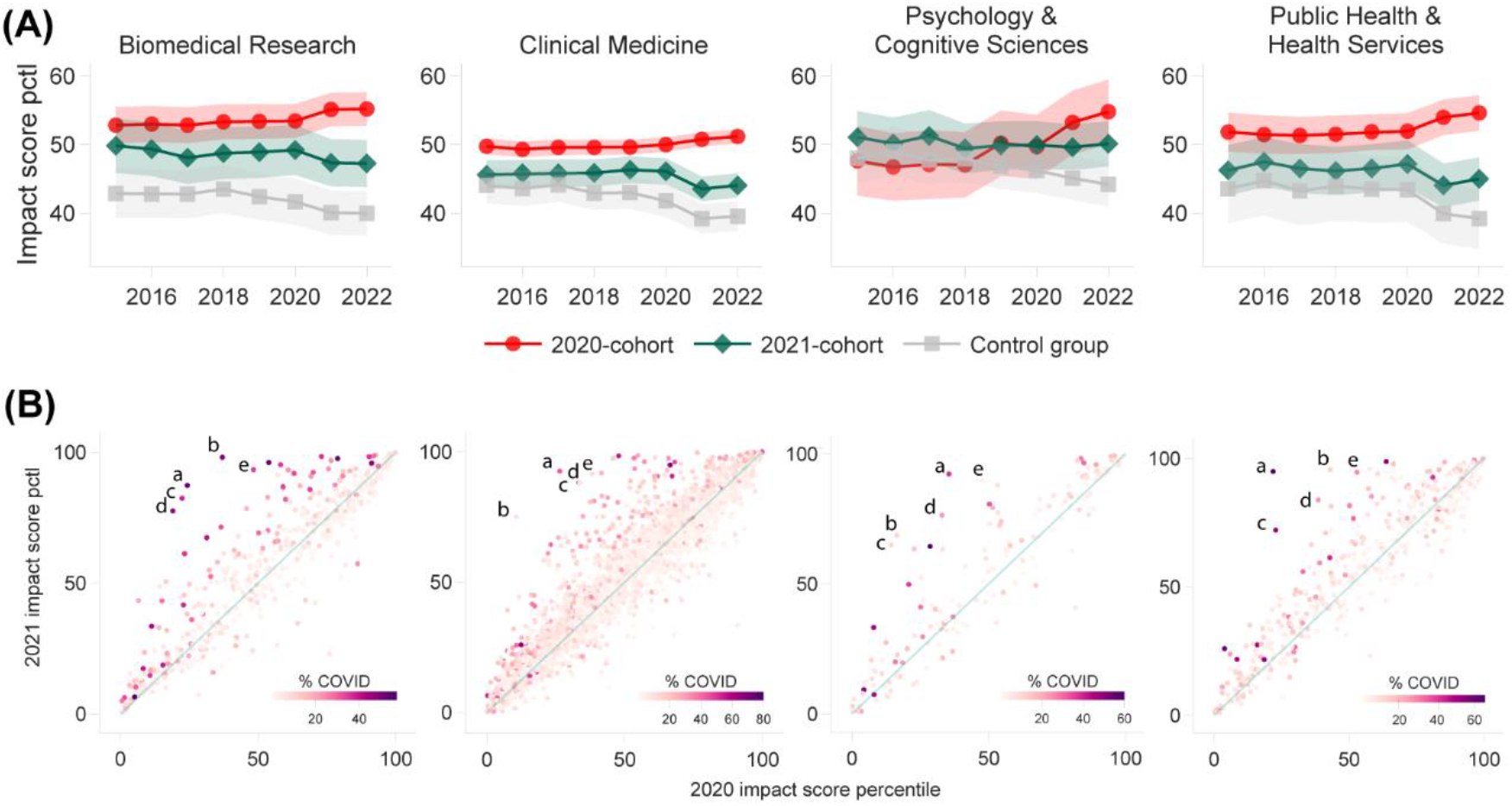
Journal rank changes based on impact score percentile. *Note.* (A) The evolution of impact score percentile from 2015-2022. The markers show the mean impact score percentiles for each year. The shaded areas show 95% confidence intervals. (B) Comparison of 2020 and 2021 impact score percentiles for 2020-cohort journals. Each point represents a journal. Points above the diagonal (shown as a light green line) have higher impact score percentiles in 2021 than in 2020. Points are colored according to the corresponding journal’s percentage of COVID-19 papers. The top 5 journals with the highest increases are labeled by *a-e* sequentially. See **Table S4** for these journals’ names.

## Discussion and conclusions

COVID-19 has been one of the most remarkable research hotspots in the recent decade. Using COVID-19 as a case, we contributed new knowledge about how research hotspots could boost the journal citation impacts by investigating the dynamic process and mechanisms of citation premium brought by COVID-19 papers. By comparing the effect of COVID-19 papers on journal citations, we found that for health sciences fields, in contrast to the initial surge of citations when COVID-19 papers began, the citation premium soon declined in 2021, leaving limited opportunities for late-entry journals to benefit from the citation boost. This finding confirms the previous hypothesis that the citation premium for late-published COVID-19 papers would be restricted as the pool of COVID-19 papers grew (Ioannidis et al., 2022). Similarly, for other research hotspots, it is also observed that following the research hotspots brings the advantage of accumulating more citations (Meyer et al., 2010). However, new publications seemed to have a decreasing scientific impact as the research field became saturated, as shown by previous examples of graphene and artificial intelligence (Klincewicz, 2016; Niu et al., 2016).

However, not all journals completely lost the citation premium brought about by COVID-19 in 2021. Albeit to a lesser extent, in certain fields, publishing COVID-19 papers in 2021 still boosted a journal’s citation impact if the journal had already published COVID-19 papers in 2020, which was not as evident for publishing COVID-19 papers published in 2021. These findings provide evidence for a “gold rush” pattern, where early entrants in an emerging and fast-evolving field are more likely to establish their advantage and gain broad and long-term benefits (Chu & Evans, 2021; Munari & Toschi, 2014). While the gold rush may also open a “window of opportunity” (Klincewicz, 2016) for late-coming journals to derive citation benefits from the topic’s popularity, our study suggests that the window of opportunity for COVID-19 is closing quickly.

Despite the inflated publication volumes in the COVID-19 period observed in Figure 1D, our data did not show increases in the citations to non-COVID-19 papers published along with COVID-19 papers in the same journal. In certain fields, non-COVID-19 papers might even receive slightly fewer citations as their journals started to publish more COVID-19 papers. In other words, the findings do not show a spillover effect over other papers of different topics for the citations boosted by research hotspots, which aligns with previous studies such as Delardas & Giannos (2022). For journals, it reflects that in a short term, the overall citation impact of a journal, as contributed by various topics, was not holistically enhanced by publishing hot topic papers and gaining temporary attention from the readers. Although previous research found that hit papers available online could increase the exposure of other papers juxtaposed on the same webpage (Kässi & Westling, 2013), it may be difficult or take a longer time to transfer the extra exposure contributed by hit papers to the entire journal’s citation increase. For researchers, the relatively disadvantaged citations to non-COVID-19 papers highlight that besides fewer opportunities to publish non-COVID-19 papers in global health sciences journals (Delardas & Giannos, 2022; He et al., 2023), researchers may also face challenges in gaining recognition and visibility amidst the overwhelming focus on pandemic-focused research. It adds additional evidence that it is problematic to evaluate researchers solely based on their publication venues without considering their individual paper topics and domains (Moed & Halevi, 2015).

The heterogeneity test shows that COVID-19 papers published in high-impact journals increased the journal and non-COVID-19 papers’ citation impact more in most fields in 2020. This finding can be associated with the evidence that prestigious journals gain more citation benefits from publishing COVID-19 papers than low-impact journals (You et al., 2023). It is possible that COVID-19 papers published in high-impact journals attracted more attention and citations than those in low-impact journals, even when the quality is similar (Larivière & Gingras, 2010). However, it should be noted that the heterogeneous effects cannot be interpreted causally because we did not control for omitted variables or reverse causality.

The findings from our analysis add a critical dimension to the ongoing debate about the appropriateness of using JIF and similar citation-based metrics as a measure of journal prestige and quality. We empirically show that the calculation methodology of JIF can be significantly influenced by highly cited papers focusing on trending topics, benefiting for a more extended period than journals that engage with these topics later. This lends support to critiques of the JIF for being susceptible to manipulation (Siler & Larivière, 2022) and steering researchers toward hot research topics popular in high-impact journals (Moustafa, 2015). This susceptibility suggests that JIF may seriously diverge from the scientific value and influence of the journal’s publications (Larivière & Gingras, 2010), potentially disadvantaging early-career and underrepresented researchers during academic evaluations (Arabi et al., 2023; Berenbaum, 2019). Consequently, this phenomenon raises questions about the equity and fairness of using JIF as a benchmark in academic evaluation. Our findings thus strengthen the argument for a more nuanced, multifaceted, and well-normalized approach to assessing journal quality and influence, moving beyond the limitations of single metrics.

In conclusion, this study suggests that journals publishing early COVID-19 papers are more likely to share shrinking but lasting citation premiums contributed by COVID-19 papers. It highlights the benefits of timely research and publishing in emerging and frontier topics, which is also supported by other empirical evidence (Huang et al., 2022). Reportedly, some journals expedited the peer review process for COVID-19 papers to swiftly communicate crucial scientific findings in response to the worldwide pandemic (McDonald et al., 2023). We concur that this accelerated peer review process, in conjunction with factors like the urgency of the COVID-19 subject matter, likely played a role in the heightened citation premium observed in our investigation. However, the reason behind the citation premium remains beyond the scope of our available data. Our analysis enables us to deduce solely that journals that released COVID-19 papers in the early stages exhibit this citation premium. Our study also cautions against using quantitative citation-based journal metrics to evaluate the quality of journals or their publications, which may be seriously affected by the citation premium of a few papers due to their topics.

As a limitation, a more expanded citation window, such as five years, may help capture a more long-term impact of the COVID-19 interruption. The thresholds and journal grouping criteria selected in this study do not necessarily represent the optimal or most universally applicable values. Additionally, evidence shows that while citations continue to grow over time, the citation differences among papers remain consistent or increase over time, with some variations among fields (Anauati et al., 2016; Galiani & Gálvez, 2017). Future studies can examine this effect again once an extended citation window is possible.

## Acknowledgments

We gratefully thank the ICSR Lab and Elsevier for sharing the Scopus data and computational resources. We appreciate Dr. Ian Hutchins for his constructive comments.

## Data Availability

Codes for this project are available on GitHub (https://github.com/UWMadisonMetaScience/covidcites). For research purposes, readers can contact Elsevier for access to Scopus data.

