## supplemental materials for "The significant yet short-term influence of research covidization on journal citation metrics"

* Chaoqun Ni

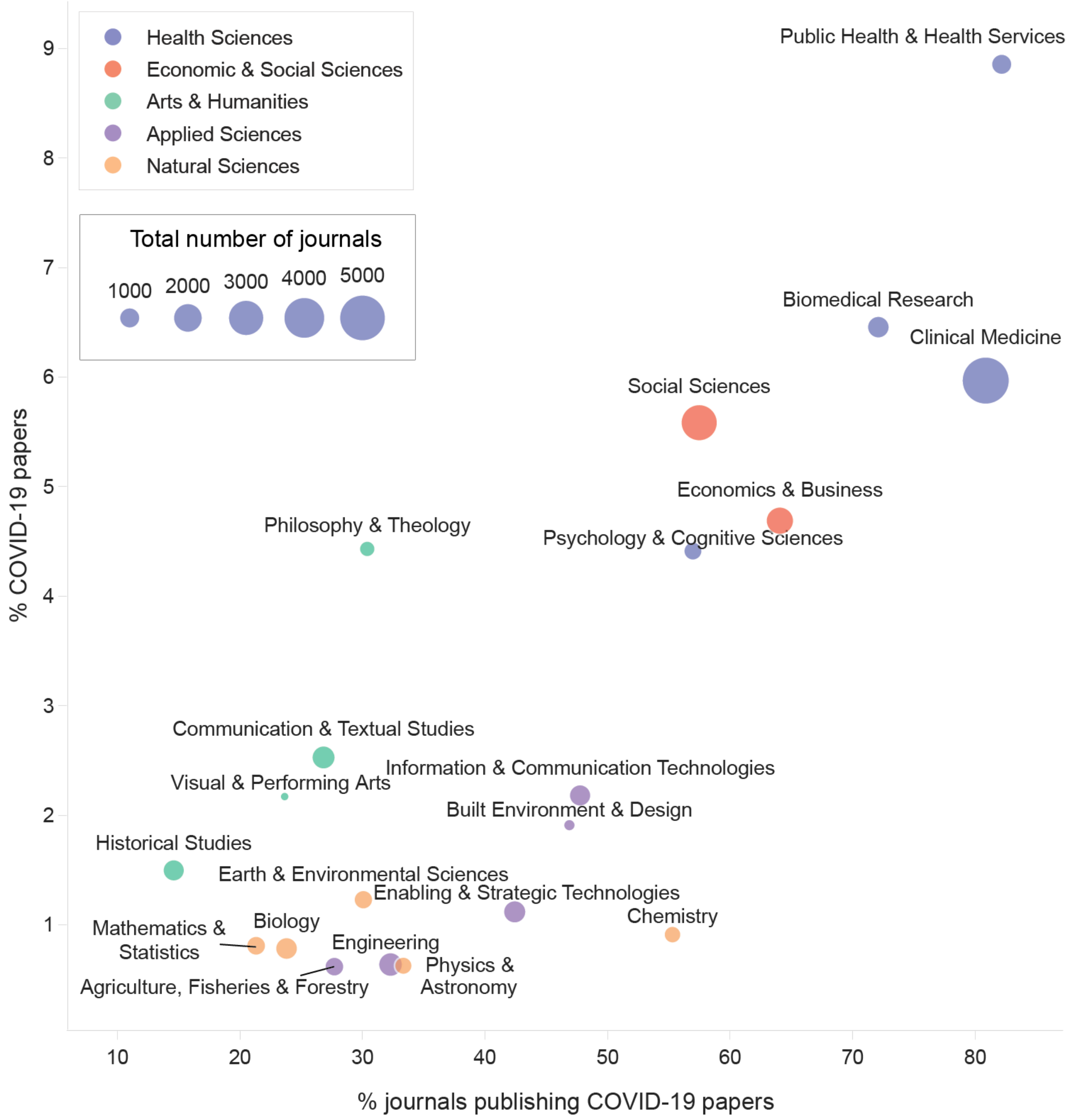


**Figure S1. Percentages of COVID-19 papers and journals publishing COVID-19 papers across fields (2020-2021)**. Node colors correspond to the domains of the fields. Node size is proportional to the total number of journals in our sample.

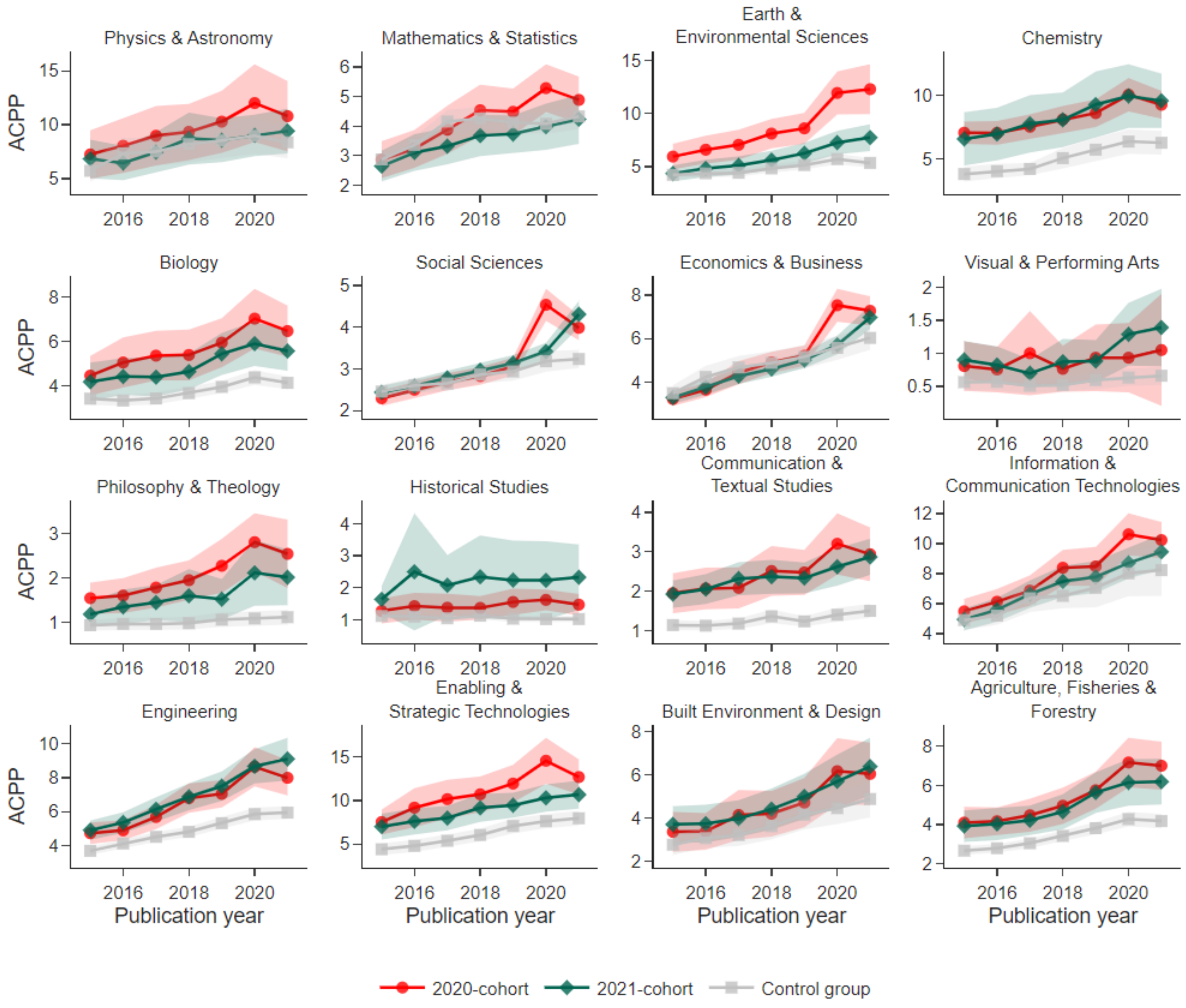


**Figure S2. The evolution of ACPP from 2015-2021 for fields outside *Health Sciences*.** The markers show the mean values of the year. The shaded areas show 95% confidence intervals.

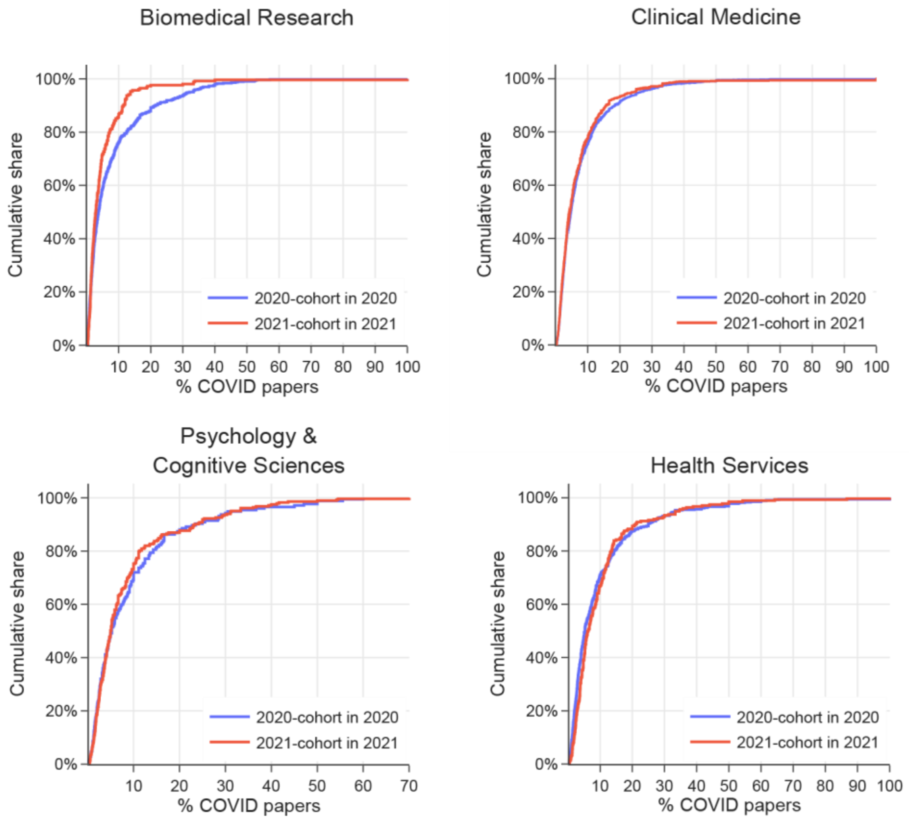


**Figure S3. COVID paper percentage cumulative distributions across fields.**

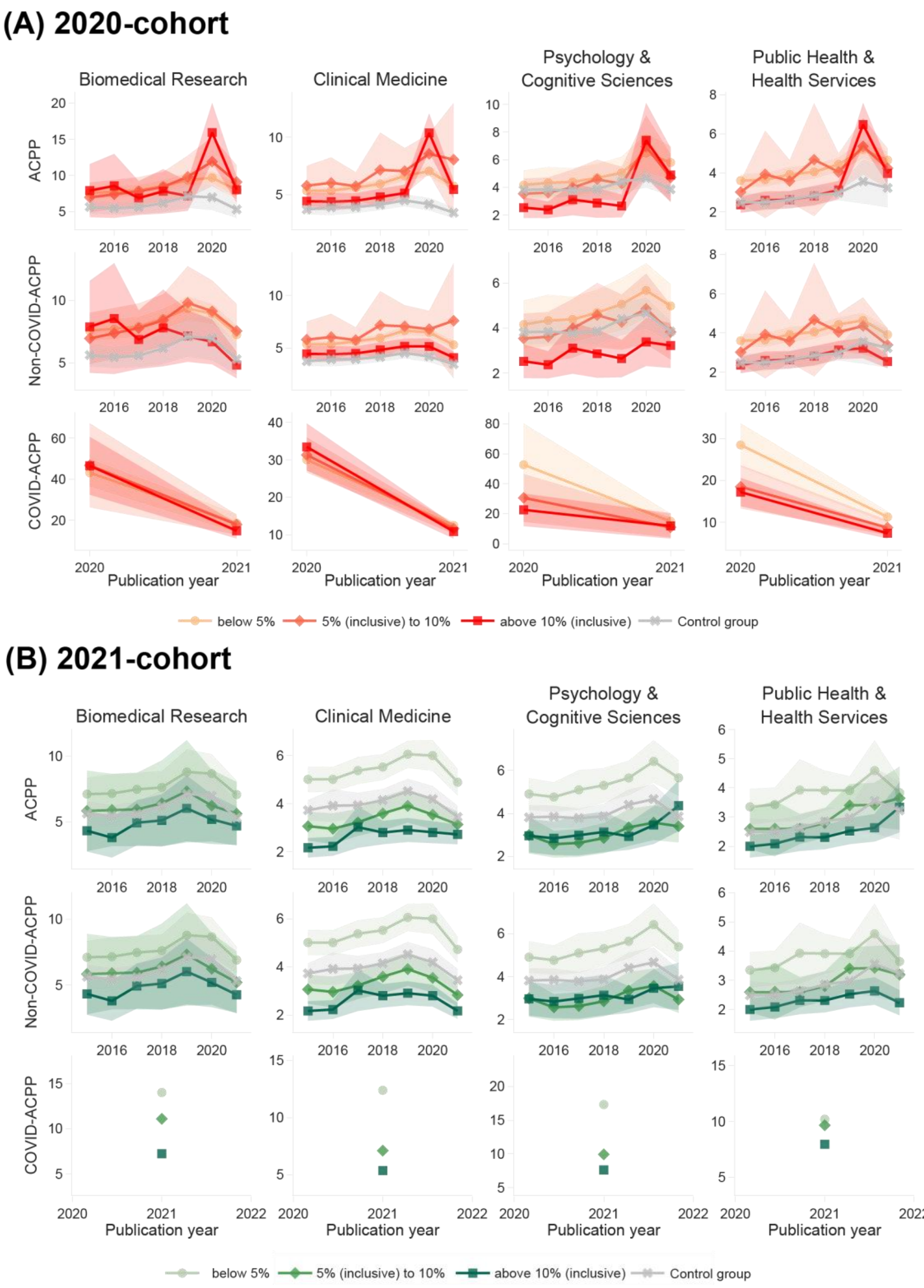


**Figure S4. The evolution of ACPP, non-COVID-ACPP, and COVID-ACPP from 2015-2021 by COVID-19 percentage.** The markers show the mean values of the year. The shaded areas show 95% confidence intervals. Journals are grouped at cutoffs of 5% and 10% of COVID-19 papers in their first year of publishing COVID-19 research (2020 for 2020-cohort, and 2021 for 2021-cohort).

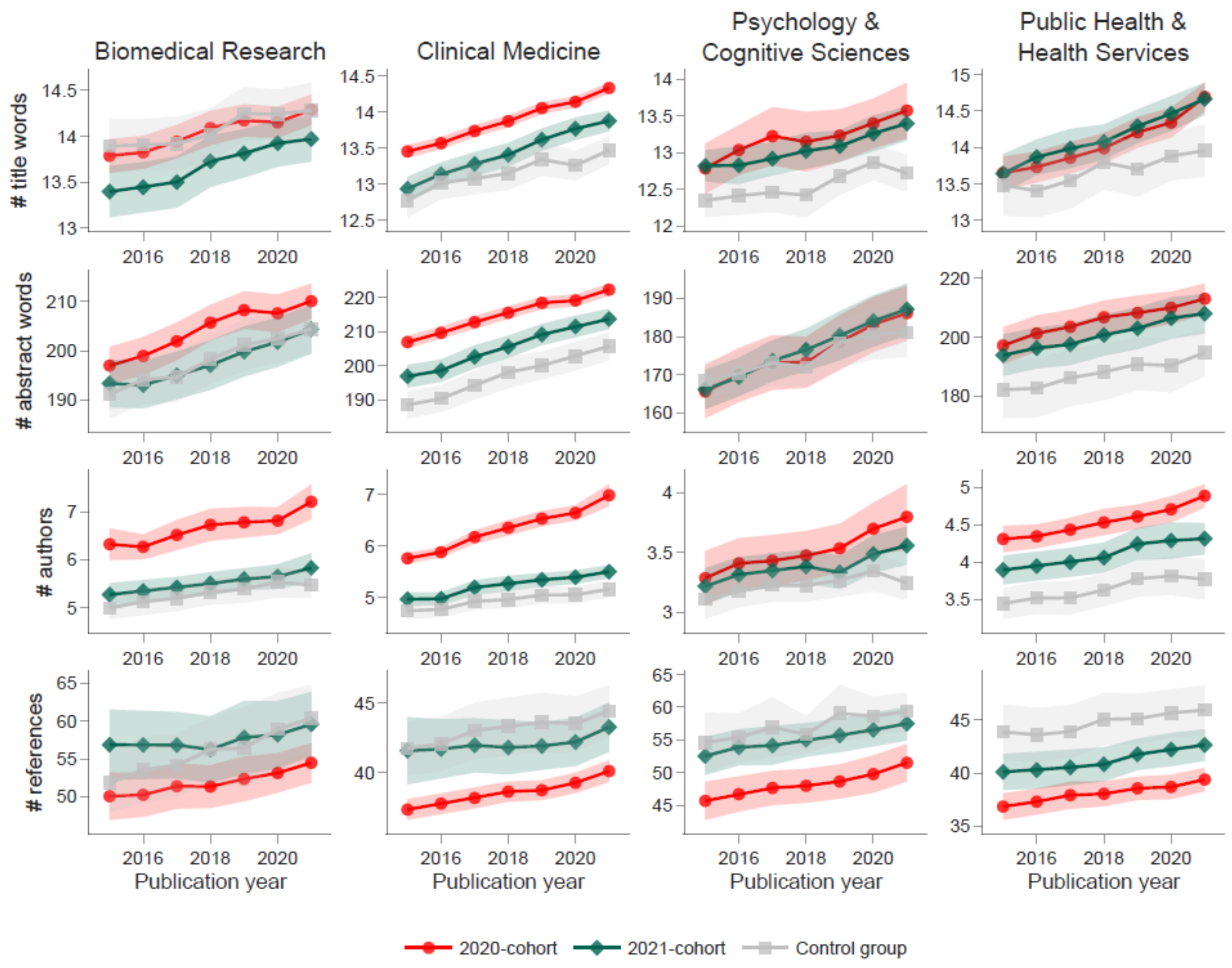


**Figure S5. The evolution of journals' average counts of title words, abstract words, authors, and references from 2015-2021.** The markers show the mean values of the year. The shaded areas show 95% confidence intervals.

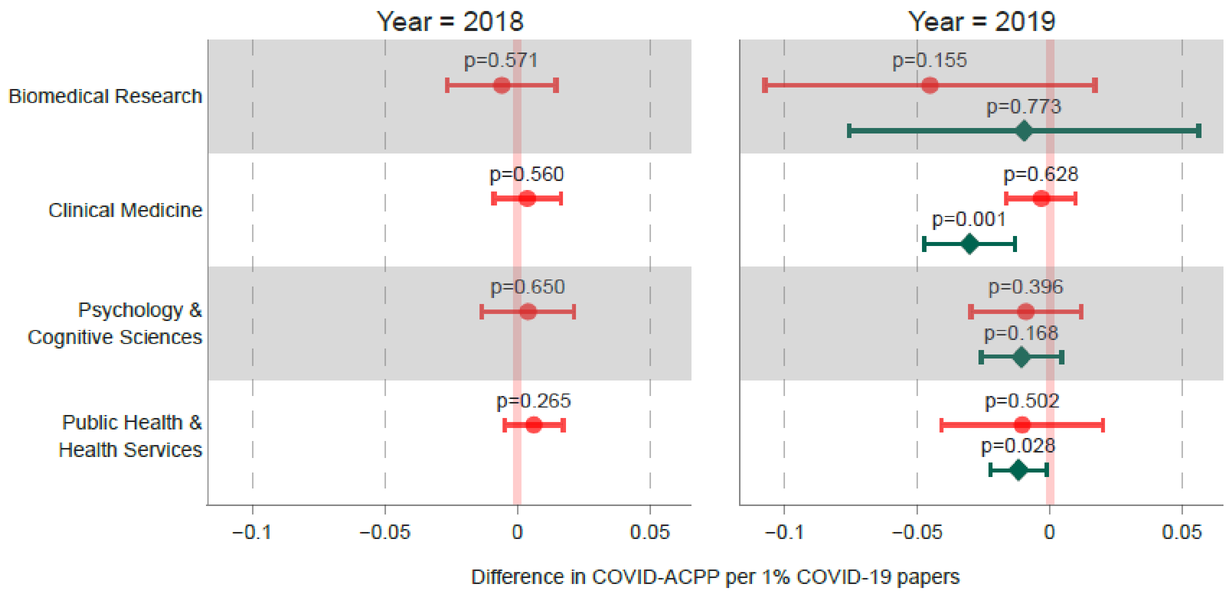


**Figure S6. Estimates of the citation effect of COVID-19 papers in placebo treatment years (2018-2019).** Estimates replicate the baseline specification. Symbols in this figure are identical to those shown in **Figure 2**.

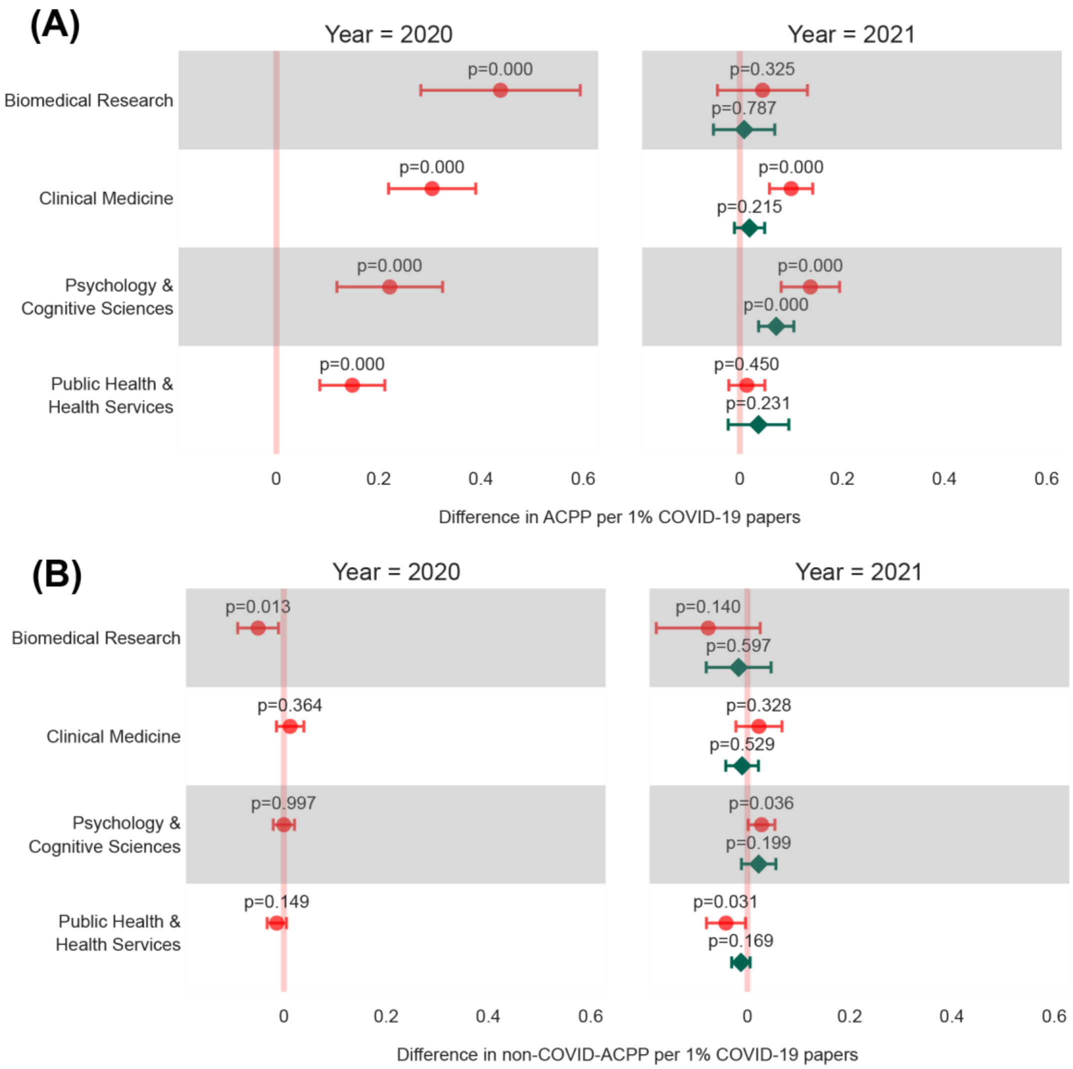


**Figure S7. Estimates of the effect of publishing COVID-19 papers using the balanced panel data.** Estimates replicate the baseline specification. Symbols in this figure are identical to those shown in **Figure 2**.

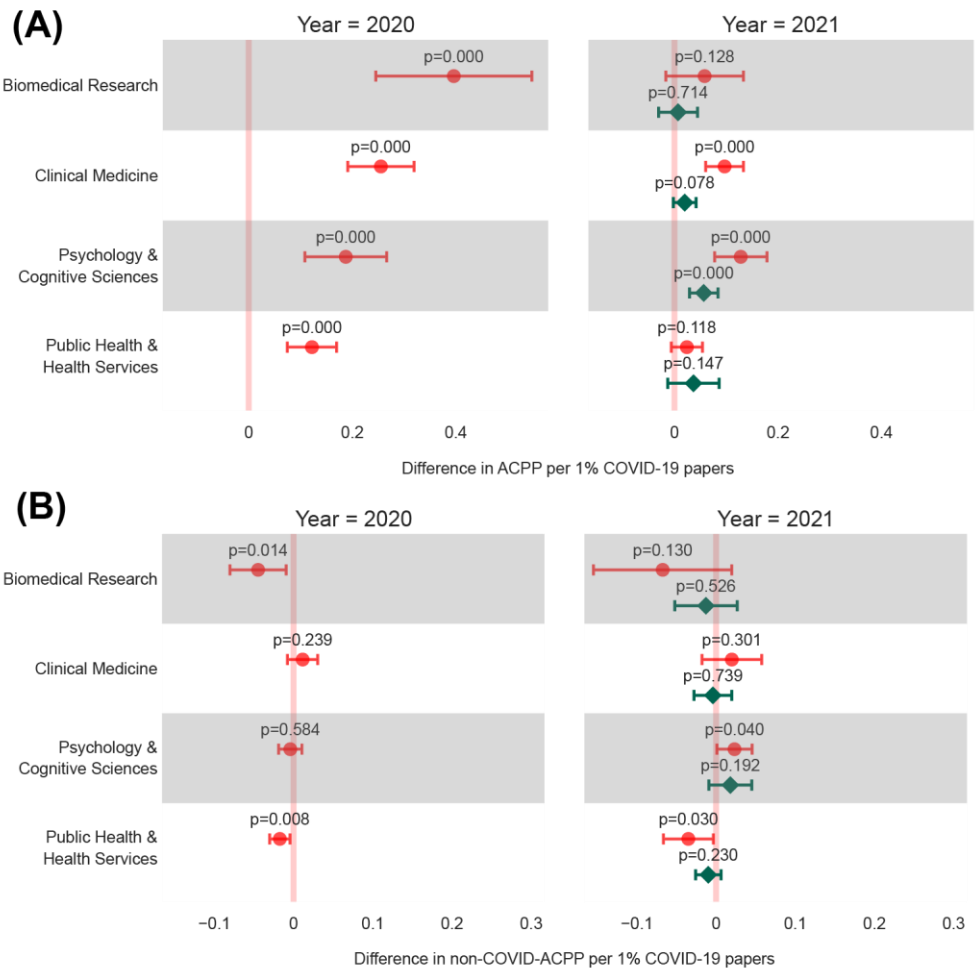


**Figure S8. Estimates of the effect of publishing COVID-19 papers using full citation counts.** Journal self-citations are included. Estimates replicate the baseline specification. Symbols in this figure are identical to those shown in **Figure 2**.

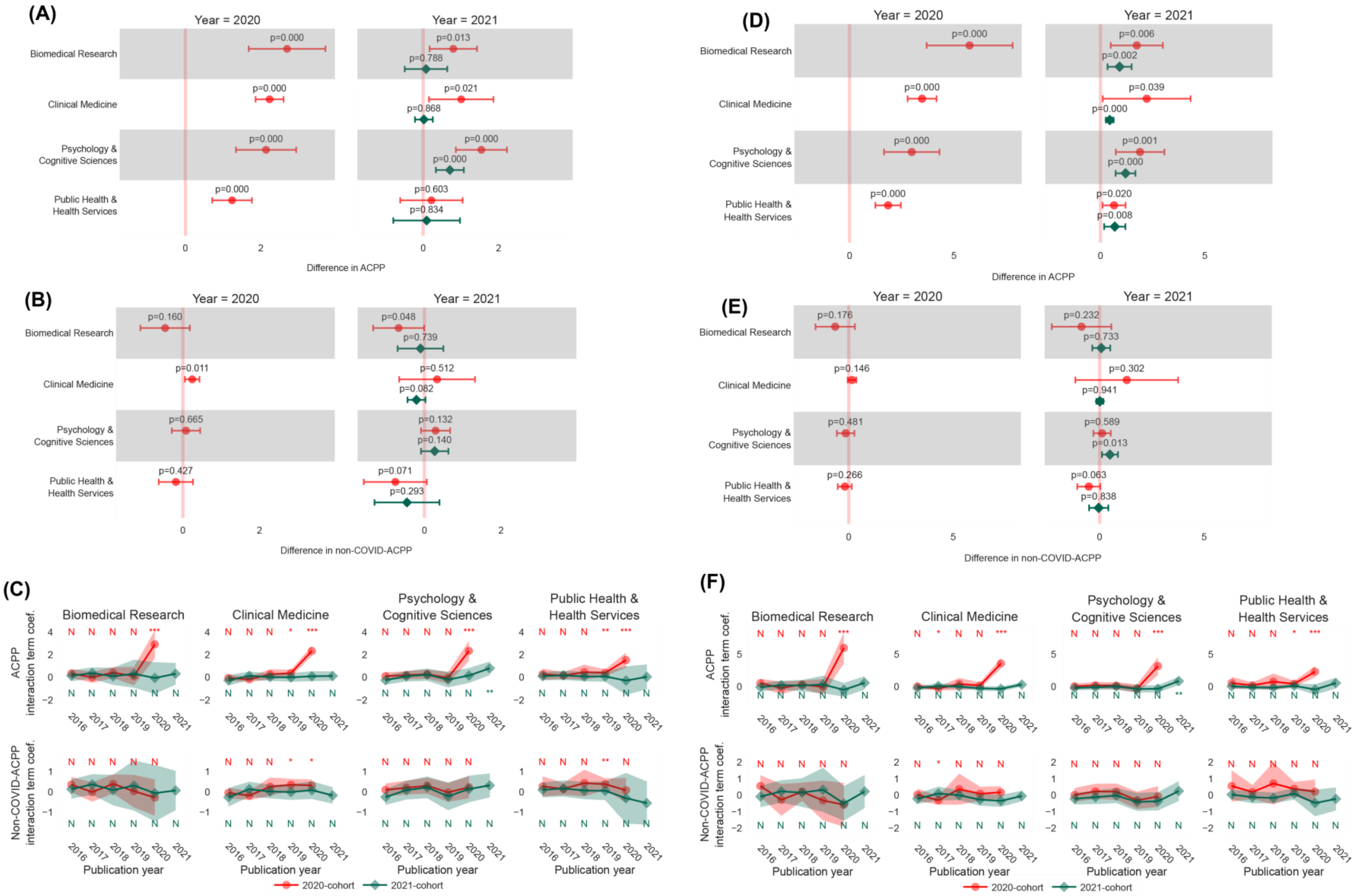


**Figure S9. Estimates of the citation effect of publishing COVID-19 papers using binary treatment variables.** The percentage of COVID-19 papers are dichotomized into a binary variable by the cutoff of 0% **(A-C)** and 5% **(D-F)**. Estimates replicate the baseline specification. Symbols in this figure are identical to those shown in **Figure 2** and **Figure 3**.

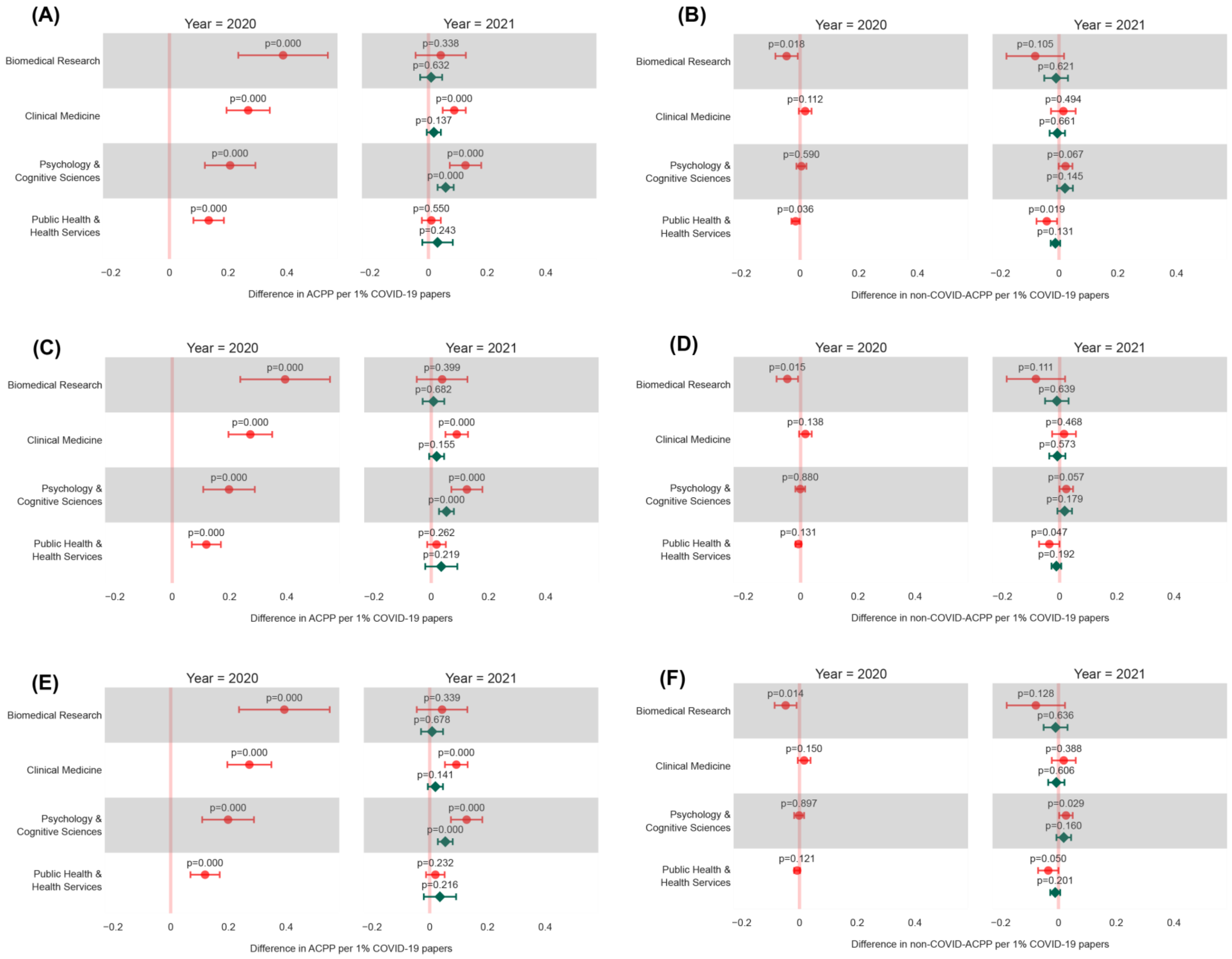


**Figure S10. Estimates of the effect of publishing COVID-19 papers controlling for journal prestige and publication volume.** Journal prestige is measured by JIF **(A-B)** and SJR **(C-D)**, SNIP **(E-F)**. Symbols in this figure are identical to those shown in **Figure 2**.

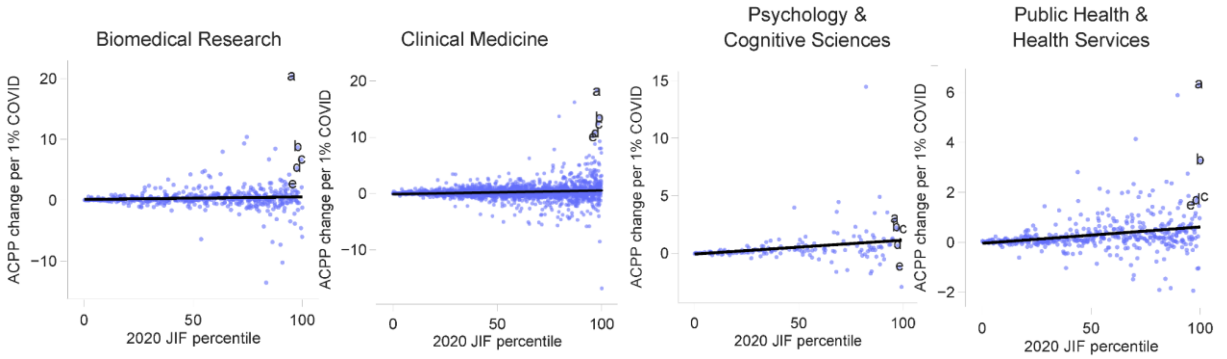


| | Biomedical Research | Clinical Medicine | Psychology & Cognitive Sciences | Public Health & Health Services |
|---|---|---|---|---|
| a | *Experimental and Molecular Medicine* | *Nature Reviews Cardiology* | *Nature Human Behavior* | *International Journal of Behavioral Nutrition and Physical Activity* |
| b | *Nature Microbiology* | *Lancet* | *Psychological Science* | *MMWR Surveillance Summaries* |
| c | *Cell* | *Nature Reviews Nephrology* | *Brain Informatics* | *Cannabis and Cannabinoid Research* |
| d | *Cell Research* | *JAMA Neurology* | *Child Development Perspectives* | *International Journal of Nursing Studies* |
| e | *Journal of Biomedical Science* | *International Journal of Oral Science* | *Neuroscience and Biobehavioral Reviews* | *Experimental Gerontology* |

**Figure S11. Correlation between the journal prestige (measured by 2020 JIF percentile) and 2020 ACPP per 1% COVID-19 papers.** Journals with the highest 2020 ACPP per 1% COVID-19 papers among top 5% journals are labeled by *a-e* sequentially.

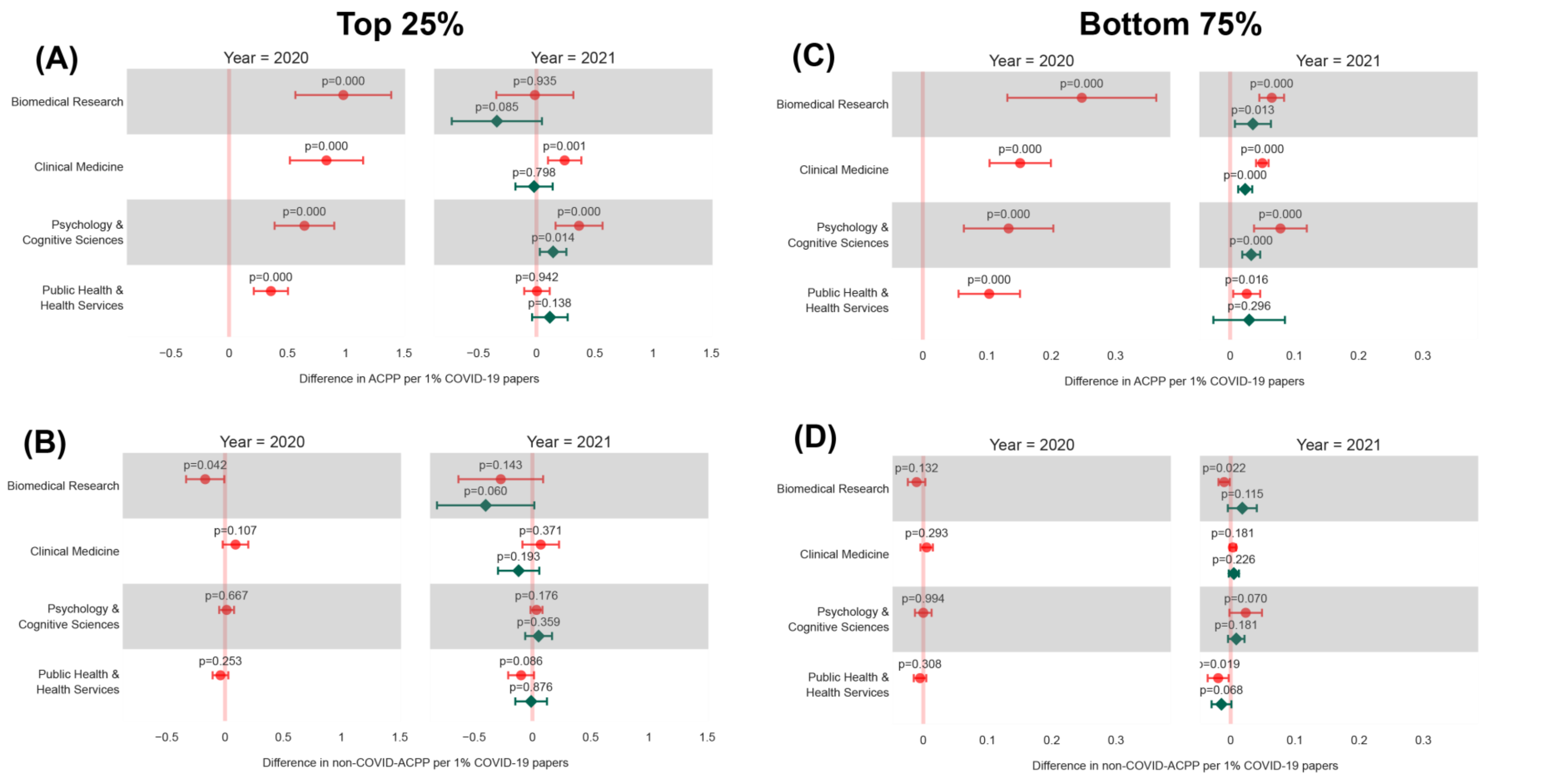


**Figure S12. Estimates of the effect of publishing COVID-19 papers by 25% high- (A-B) and 75% low-prestige (C-D) journals.** Estimates replicate the baseline specification. Symbols in this figure are identical to those shown in **Figure 2**.

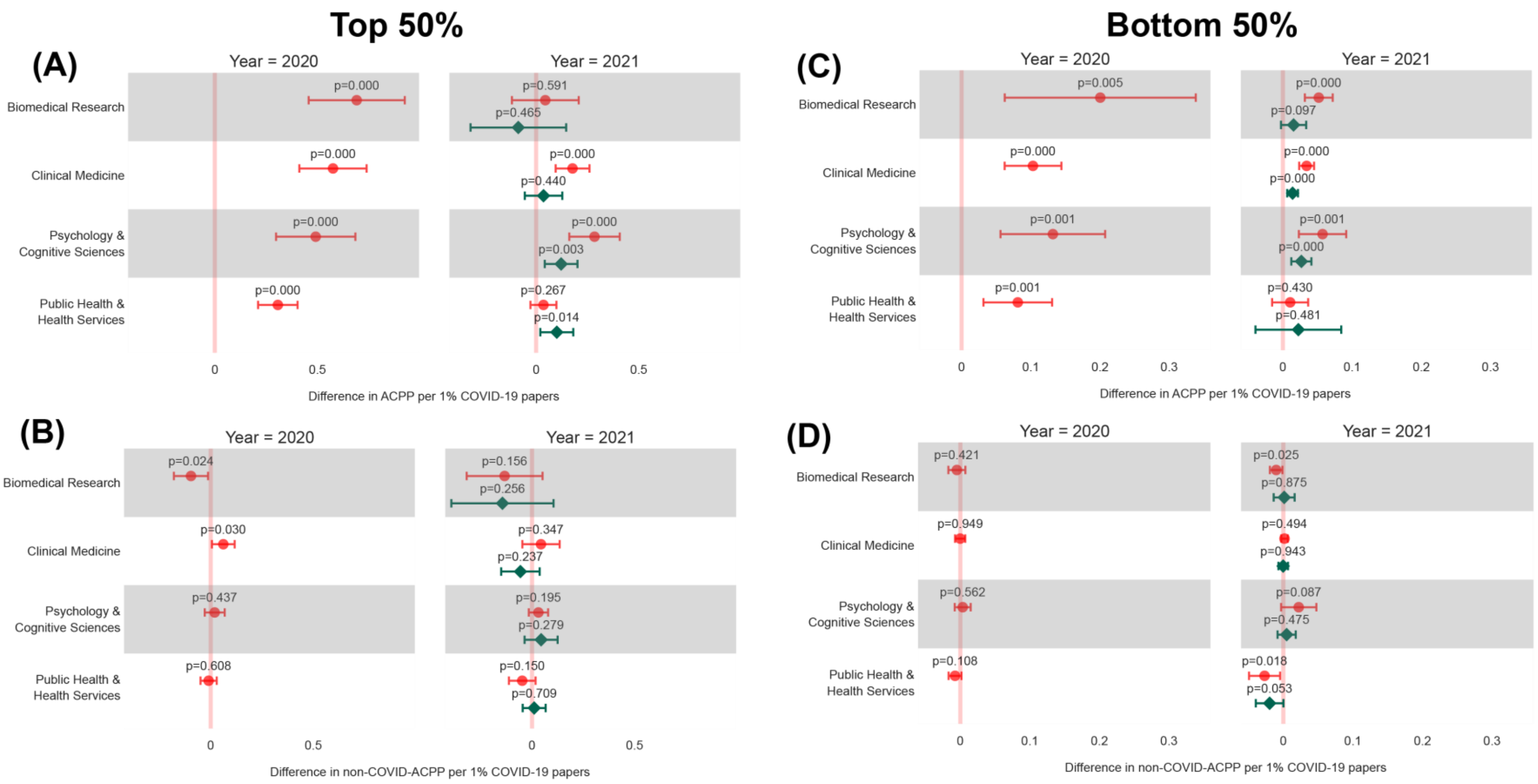


**Figure S13. Estimates of the effect of publishing COVID-19 papers by 50% high- (A-B) and 50% low-prestige (C-D) journals.** Estimates replicate the baseline specification. Symbols in this figure are identical to those shown in **Figure 2**.

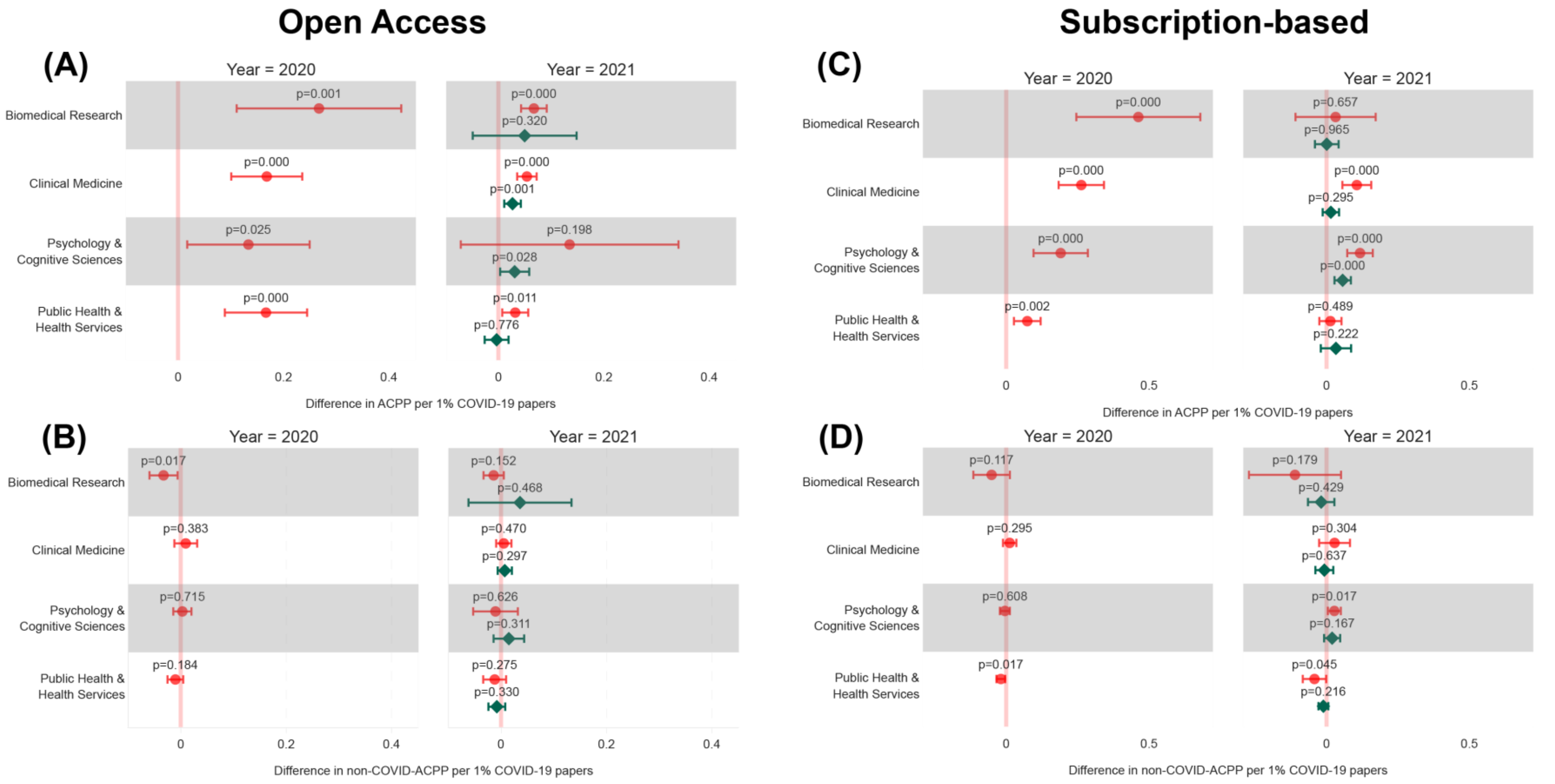


**Figure S14. Estimates of the effect of publishing COVID-19 papers by OA (A-B) and subscription-based (C-D) journals.** Estimates replicate the baseline specification. Symbols in this figure are identical to those shown in **Figure 2**.

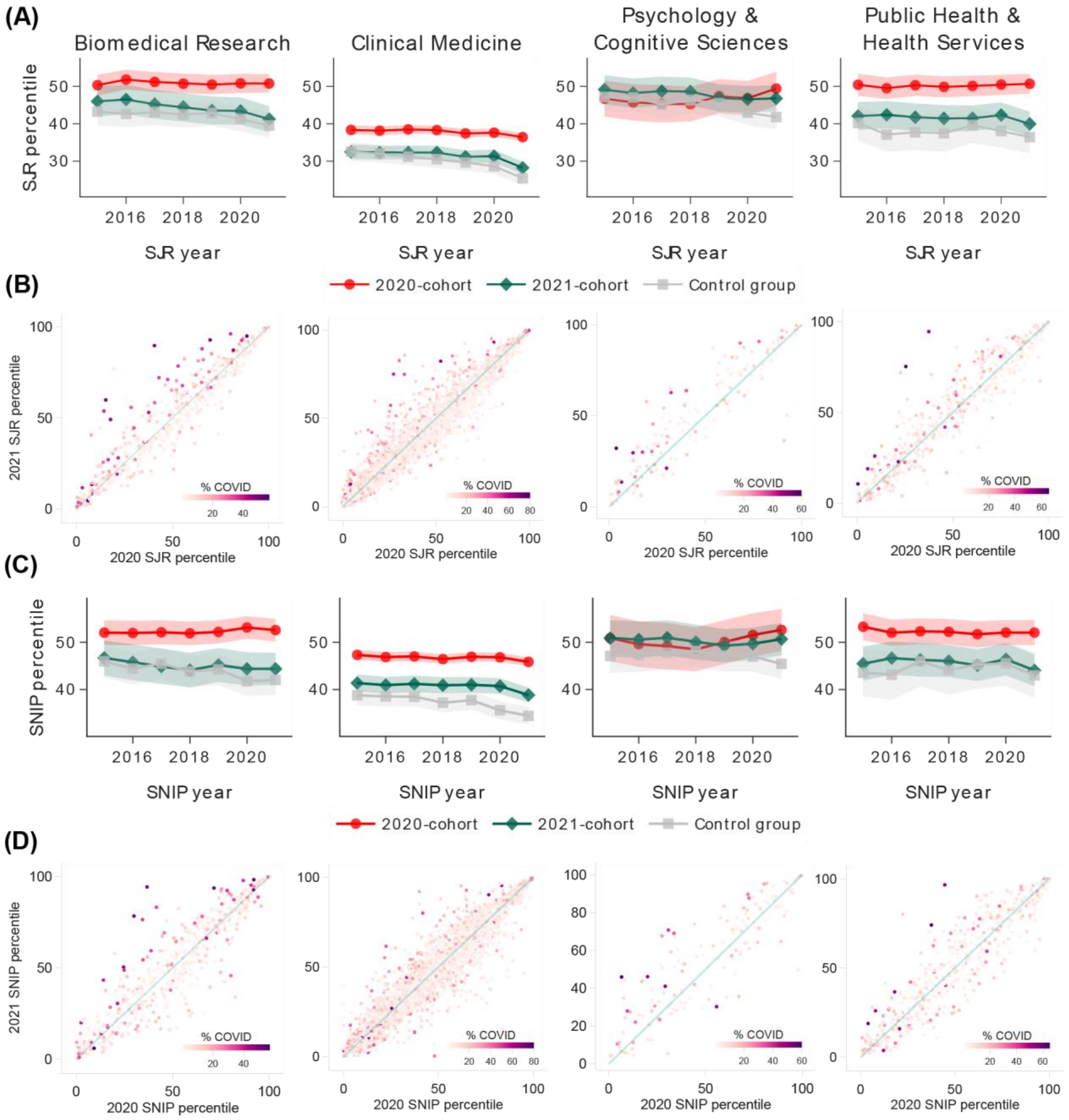


**Figure S15. Journal rank changes based on SJR (A-B) and SNIP (C-D).** SJR and SNIP for 2022 were not available by the date of data curation. Symbols in this figure are identical to those shown in **Figure 1.**

**Table S1. Top 5 journals with the highest increase in ACPP from 2019 to 2020.** Journals are ordered by their increase in ACPP.

| | Journal name | 2019 ACPP | 2020 ACPP | Increase in ACPP | % COVID-19 papers |
|---|---|---|---|---|---|
| | ***Biomedical Research*** | | | | |
| a | *Cell* | 88.06 | 187.10 | 99.04 | 14.91 |
| b | *Lancet Infectious Diseases* | 38.86 | 123.50 | 84.64 | 28.34 |
| c | *Journal of Infection* | 12.03 | 88.76 | 76.74 | 36.36 |
| d | *Journal of Medical Virology* | 4.91 | 73.81 | 68.90 | 48.88 |
| e | *Nature Microbiology* | 39.32 | 105.53 | 66.21 | 7.63 |
| | ***Clinical Medicine*** | | | | |
| a | *Lancet* | 102.40 | 363.43 | 261.03 | 19.35 |
| b | *New England Journal of Medicine* | 105.35 | 254.97 | 149.62 | 18.65 |
| c | *Lancet Respiratory Medicine* | 53.60 | 183.32 | 129.72 | 32.84 |
| d | *Lancet Public Health* | 35.73 | 149.73 | 113.99 | 27.45 |
| e | *JAMA* | 30.26 | 120.70 | 90.44 | 14.08 |
| | ***Psychology & Cognitive Sciences*** | | | | |
| a | *Nature Human Behavior* | 12.91 | 60.23 | 47.32 | 15.97 |
| b | *British Journal of Health Psychology* | 7.59 | 25.80 | 18.21 | 28.33 |
| c | *Journal of Loss and Trauma* | 2.20 | 20.29 | 18.09 | 38.46 |
| d | *Journal of the American Academy of Child and A...* | 13.89 | 31.98 | 18.08 | 1.25 |
| e | *Journal of Anxiety Disorders* | 12.47 | 30.40 | 17.93 | 14.85 |
| | ***Public Health & Health Services*** | | | | |
| a | *MMWR Surveillance Summaries* | 51.50 | 82.62 | 31.12 | 9.52 |
| b | *JMIR Public Health and Surveillance* | 6.00 | 35.44 | 29.44 | 61.62 |
| c | *Epidemiology and health* | 7.02 | 33.25 | 26.23 | 28.13 |
| d | *European Journal of Epidemiology* | 16.23 | 41.73 | 25.50 | 17.53 |
| e | *Journal of Aging and Social Policy* | 3.21 | 25.20 | 21.99 | 54.90 |

**Table S2. Results of the in-space placebo test.**

| Outcome | Cohort | Year | Empirical p-values | | | |
|---|---|---|---|---|---|---|
| | | | ***Biomedical Research*** | ***Clinical Medicine*** | ***Psychology & Cognitive Sciences*** | ***Public Health & Health Services*** |
| ACPP | 2020-cohort | 2020 | 0.000 | 0.000 | 0.000 | 0.000 |
| | 2020-cohort | 2021 | 0.000 | 0.004 | 0.000 | 0.002 |
| | 2021-cohort | 2021 | 0.840 | 0.356 | 0.046 | 0.086 |
| Non-COVID-ACPP | 2020-cohort | 2020 | 0.000 | 0.000 | 0.438 | 0.008 |
| | 2020-cohort | 2021 | 0.166 | 0.132 | 0.594 | 0.000 |
| | 2021-cohort | 2021 | 0.384 | 0.856 | 0.910 | 0.052 |

**Table S3. Heterogeneity analysis of COVID-19 research effects on ACPP by journal prestige and OA status**

| Field | Journal cohort | Publication year | Coef. | p-value | Confidence interval | |
|---|---|---|---|---|---|---|
| | | | | | Lower | Upper |
| ***25% high vs. 75% low-prestige*** | | | | | | |
| Biomedical Research | 2020 | 2020 | 0.802 | 0.000 | 0.402 | 1.201 |
| | 2020 | 2021 | -0.068 | 0.627 | -0.345 | 0.208 |
| | 2021 | 2021 | -0.360 | 0.032 | -0.689 | -0.030 |
| Clinical Medicine | 2020 | 2020 | 0.741 | 0.000 | 0.475 | 1.008 |
| | 2020 | 2021 | 0.220 | 0.046 | 0.004 | 0.435 |
| | 2021 | 2021 | -0.012 | 0.725 | -0.078 | 0.054 |
| Psychology & Cognitive Sciences | 2020 | 2020 | 0.582 | 0.000 | 0.321 | 0.844 |
| | 2020 | 2021 | 0.268 | 0.008 | 0.072 | 0.463 |
| | 2021 | 2021 | 0.089 | 0.095 | -0.015 | 0.193 |
| Public Health & Health Services | 2020 | 2020 | 0.302 | 0.000 | 0.181 | 0.424 |
| | 2020 | 2021 | -0.023 | 0.568 | -0.102 | 0.056 |
| | 2021 | 2021 | 0.074 | 0.252 | -0.053 | 0.200 |
| ***50% high vs. 50% low-prestige*** | | | | | | |
| Biomedical Research | 2020 | 2020 | 0.537 | 0.000 | 0.285 | 0.789 |
| | 2020 | 2021 | -0.022 | 0.759 | -0.162 | 0.118 |
| | 2021 | 2021 | -0.150 | 0.176 | -0.368 | 0.068 |
| Clinical Medicine | 2020 | 2020 | 0.507 | 0.000 | 0.360 | 0.655 |
| | 2020 | 2021 | 0.138 | 0.020 | 0.022 | 0.254 |
| | 2021 | 2021 | 0.009 | 0.747 | -0.047 | 0.066 |
| Psychology & Cognitive Sciences | 2020 | 2020 | 0.417 | 0.000 | 0.215 | 0.619 |
| | 2020 | 2021 | 0.212 | 0.001 | 0.089 | 0.336 |
| | 2021 | 2021 | 0.076 | 0.044 | 0.002 | 0.150 |
| Public Health & Health Services | 2020 | 2020 | 0.260 | 0.000 | 0.170 | 0.350 |
| | 2020 | 2021 | 0.021 | 0.384 | -0.027 | 0.070 |
| | 2021 | 2021 | 0.068 | 0.117 | -0.017 | 0.154 |
| ***OA vs. subscription-based*** | | | | | | |
| Biomedical Research | 2020 | 2020 | -0.180 | 0.153 | -0.427 | 0.067 |
| | 2020 | 2021 | 0.039 | 0.549 | -0.089 | 0.167 |
| | 2021 | 2021 | 0.055 | 0.274 | -0.044 | 0.154 |
| Clinical Medicine | 2020 | 2020 | -0.101 | 0.033 | -0.195 | -0.008 |
| | 2020 | 2021 | -0.060 | 0.110 | -0.134 | 0.014 |
| | 2021 | 2021 | 0.006 | 0.531 | -0.012 | 0.024 |
| Psychology & Cognitive Sciences | 2020 | 2020 | -0.067 | 0.364 | -0.211 | 0.078 |
| | 2020 | 2021 | 0.032 | 0.752 | -0.168 | 0.232 |
| | 2021 | 2021 | -0.011 | 0.542 | -0.046 | 0.024 |
| Public Health & Health Services | 2020 | 2020 | 0.084 | 0.039 | 0.004 | 0.164 |
| | 2020 | 2021 | 0.008 | 0.638 | -0.026 | 0.043 |
| | 2021 | 2021 | -0.048 | 0.056 | -0.098 | 0.001 |

**Table S4. Top 5 journals with the highest increases in JIF percentile.** Letters correspond to the labels in **Figure 4B**.

| | ***Biomedical Research*** | ***Clinical Medicine*** | ***Psychology & Cognitive Sciences*** | ***Public Health & Health Services*** |
|---|---|---|---|---|
| a | *Osong Public Health and Research Perspectives* | *Clinical Neuropsychiatry* | *Journal of Loss and Trauma* | *World Medical and Health Policy* |
| b | *Journal of Medical Virology* | *Chemical Biology Letters* | *Revista de Psicopatologia y Psicologia Clinica* | *Family Medicine and Community Health* |
| c | *Current Tropical Medicine Reports* | *Irish Journal of Psychological Medicine* | *Clinica y Salud* | *Journal of Health Management* |
| d | *Infezioni in Medicina* | *GMS German Medical Science* | *Group Dynamics* | *Health Sociology Review* |
| e | *Microbes and Infection* | *Asian Journal of Psychiatry* | *Death Studies* | *Public Health Research and Practice* |
